# mRNA therapy restores ureagenesis and corrects glutathione metabolism in argininosuccinic aciduria

**DOI:** 10.1101/2022.10.19.512931

**Authors:** Sonam Gurung, Oskar V. Timmermand, Dany Perocheau, Ana Luisa Gil-Martinez, Magdalena Minnion, Loukia Touramanidou, Sherry Fang, Martina Messina, Youssef Khalil, Abigail R. Barber, Richard S. Edwards, Patrick F. Finn, Alex Cavedon, Summar Siddiqui, Lisa Rice, Paolo G.V. Martini, Philippa B. Mills, Simon N. Waddington, Paul Gissen, Simon Eaton, Mina Ryten, Martin Feelisch, Andrea Frassetto, Timothy H. Witney, Julien Baruteau

## Abstract

Argininosuccinate lyase (ASL) is a key enzyme integral to the hepatic urea cycle which is required for ammonia detoxification, and the citrulline-nitric oxide (NO) cycle for NO production. ASL deficient patients present with argininosuccinic aciduria (ASA), an inherited metabolic disease with hyperammonaemia and a chronic systemic phenotype with neurocognitive impairment and chronic liver disease. ASL deficiency as an inherited model of systemic NO deficiency, shows enhanced nitrosative and oxidative stress. Here, we describe the dysregulation of glutathione biosynthesis and upstream cysteine utilization in ASL-deficient patients and mice using targeted metabolomics and *in vivo* positron emission tomography (PET) imaging using (*S*)-4-(3-^18^F-fluoropropyl)-L-glutamate ([^18^F]FSPG). Upregulation of cysteine metabolism contrasted with glutathione depletion and down-regulated antioxidant pathways. *hASL* mRNA encapsulated in lipid nanoparticles corrected and rescued the neonatal and adult Asl-deficient mouse phenotypes, respectively, enhancing ureagenesis and glutathione metabolism and ameliorating chronic liver disease. We further present [^18^F]FSPG PET as a novel non-invasive diagnostic tool to assess liver disease and therapeutic efficacy in ASA. These findings support clinical translation of mRNA therapy for ASA.

## Introduction

Urea cycle defects (UCDs) are inborn errors of metabolism causing dysfunction in ammonia detoxification and endogenous arginine synthesis. Argininosuccinic aciduria (ASA) (OMIM 207900) is the second most common UCD, accounting for ∼16% of all UCDs (1). ASA is caused by deficiency in argininosuccinate lyase (ASL), a cytosolic urea cycle enzyme which catalyses the conversion of argininosuccinate into arginine and fumarate, thereby enabling the removal of excess nitrogen (2, 3). ASL is also involved in the citrulline-nitric oxide (NO) cycle to produce NO through the channelling of extracellular L-arginine to nitric oxide synthase (NOS) (4, 5). Early-onset patients display hyperammonemia, while late-onset patients present with either acute hyperammonaemia and/or chronic phenotype of neurocognitive impairment and liver disease (3). Compared to other UCDs, ASA is associated with a high burden of chronic liver complications such as elevated levels of serum transaminases, hepatomegaly, fibrosis, steatosis, hepatocellular injury and glycogen storage both in patients (6-10) and in the mouse model *Asl*^*Neo/Neo*^ (8, 11). Some patients present with liver cirrhosis or, rarely, hepatocellular carcinoma (9, 12, 13). The underlying processes that trigger liver disease are unclear, but suggested mechanisms include hyperammonaemia, arginine deficiency, argininosuccinate toxicity, NO deficiency and oxidative stress (6, 9). More detailed mechanistic insight into liver pathophysiology will be crucial to identifying optimal diagnostic markers for better assessment of disease severity, prediction of disease progression and assessment of response to therapy.

Current therapeutic guidelines for ASA aim to normalise ammonia and arginine levels through low protein diet, ammonia scavenger drugs and arginine supplementation. In severe cases, patients require liver transplantation. Other experimental therapies include antioxidants, autophagy enhancers, creatinine supplementation and gene therapies (2, 8, 11, 14-18). However, long-term outcome remains poor, with persisting chronic liver disease and unpredictable risk of complication towards fibrosis, cirrhosis and malignancy asserting the need for long-term liver monitoring and novel therapeutic strategies that can target the liver pathology (2).

In this study, we describe the dysregulation of liver glutathione metabolism in ASA and present positron emission tomography (PET) using the radiotracer (*S*)-4-(3-^18^F-fluoropropyl)-L-glutamate ([^18^F]FSPG) as a non-invasive diagnostic tool to assess the extent of liver disease in ASA. [^18^F]FSPG reports on the activity of the amino acid antiporter, system X_C_^-^, which under normal physiological conditions imports cystine from the blood in exchange for intracellular glutamate. Cystine is subsequently reduced to cysteine for *de novo* glutathione biosynthesis (19). Owing to the overexpression of system X_C_^-^ in malignancy, [^18^F]FSPG has shown utility in clinical trials for cancer diagnosis (20-23), and preclinically [^18^F]FSPG has been used to non-invasively assess redox status both in tumors (24-26) and in a mouse model of multiple sclerosis (27).

We further show that mRNA therapy can treat neonatal and rescue adult *Asl*^*Neo/Neo*^ mice by correcting both ureagenesis and glutathione metabolism *in vivo*, thereby demonstrating this strategy as a promising novel therapy for ASA.

## Results

### ASL-deficient patients and mouse model *Asl*^*Neo/Neo*^ show downregulation of glutathione biosynthesis despite limited evidence of oxidative stress

Previous publications have highlighted the role of oxidative stress in the pathophysiology of ASL deficiency (5, 16, 28). Therefore, we aimed to study glutathione as a key intracellular antioxidant and its metabolism in ASA, including its close interaction with the transsulfuration pathway (Figure 1A). We compared the plasma concentrations of total homocysteine, glycine and glutamate in patients from Great Ormond Street Hospital for Children, London, UK affected by one of the 3 main urea cycle defects: ornithine transcarbamylase deficiency (OTCD) (n=13), argininosuccinate synthase deficiency (ASSD) (n=10) and argininosuccinate lyase deficiency (ASLD) (n=13), excluding OTC carriers and liver-transplanted patients. Compared to OTCD and ASSD, patients with ASLD had significantly higher mean plasma levels of the 3 precursor metabolites of glutathione biosynthesis, homocysteine (Figure 1B), glutamate (Figure 1C) and glycine (Figure 1D) levels. Since the initiation of follow-up, the reports of every single measurement of plasma total homocysteine (Supplementary Figure 1A), glycine (Supplementary Figure 1B) and glutamate (Supplementary Figure 1C) levels measured in these patients confirmed the significantly increased values in ASLD compared to OTCD and ASSD. Plasma total homocysteine did not differ between early- and late-onset ASA patients (Supplementary Figure 1D).

**Figure 1.**
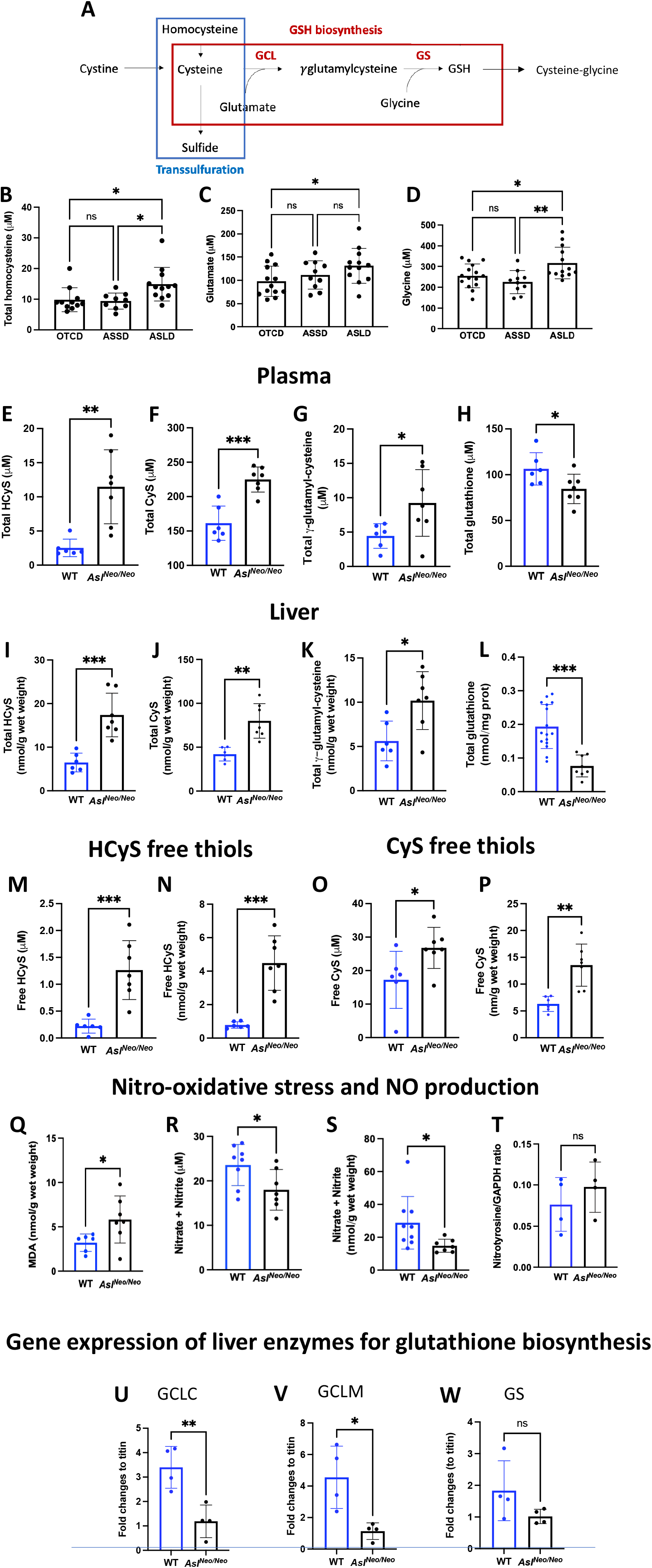
ASL-deficient patients and mouse model *Asl*^*Neo/Neo*^ show dysfunction of glutathione metabolism despite limited evidence of oxidative stress. (**A**) Glutathione biosynthesis requires precursor metabolites glutamate, glycine and cysteine, the latter being an intermediary metabolite from the transsulfuration pathway. (**B**) Mean of plasma total homocysteine, (**C**) glycine and (**D**) glutamate in patients with OTCD, ASSD and ASLD. Plasma (**E**) total homocysteine, (**F**) cysteine, (**G**) γ-glutamyl-cysteine and (**H**) total glutathione levels, Liver (**I**) homocysteine, (**J**) cysteine, (**K**) γ-glutamyl-cysteine total thiols and (**L**) total glutathione levels in 2-weeks old *Asl*^*Neo/Neo*^ mice and WT littermates. Free homocysteine in (**M**) plasma and (**N**) liver, and free cysteine in (**O**) plasma and (**P**) liver in *Asl*^*Neo/Neo*^ mice compared to WT littermates. (**Q**) Lipid peroxidation measured by thiobarbituric acid reactive substances in *Asl*^*Neo/Neo*^ mice and WT littermates. Nitric oxide metabolites (nitrite and nitrate) in (**R**) plasma and (**S**) liver samples of *Asl*^*Neo/Neo*^ mice and WT littermates. (**T**) Quantification of liver nitrotyrosine levels by western blot between *Asl*^*Neo/Neo*^ mice and WT. (**B-D**) One-way ANOVA with Tuckey’s post-test (**E-S; T**): Unpaired 2-tailed Student’s t test; * p<0.05, ** p<0.01, *** p<0.001, ns not significant. **(B-D)**: OTCD n=11-13, ASSD n=10, ASLD n=13. (**E-S**) WT n=6-8; *Asl*^*Neo/Neo*^ n=7. ASSD: argininosuccinate synthase deficiency; ASLD: argininosuccinate lyase deficiency; CyS: cysteine; GSH: glutathione; HcyS: homocysteine; MDA: malondialdehyde; OTCD: ornithine transcarbamylase deficiency. Graphs show means +SD.

The hypomorphic *Asl*^*Neo/Neo*^ mouse model recapitulates much of the human phenotype of ASLD (4, 16). Corroborating human data, total homocysteine levels in 2 week-old *Asl*^*Neo/Neo*^ mice were significantly elevated compared to WT littermates (*p*=0.002, Figure 1E). Other contributors to the glutathione biosynthesis pathway, including cysteine (Figure 1F) and γ-glutamyl-cysteine (Figure 1G), were significantly increased in *Asl*^*Neo/Neo*^ mice compared to WT. In contrast, plasma total glutathione was significantly decreased (Figure 1H). Similarly, tissue concentrations of homocysteine (Figure 1I), cysteine (Figure 1J), γ-glutamyl-cysteine (Figure 1K) and cysteine-glycine (Supplementary Figure 1E) in liver showed a significant increase in *Asl*^*Neo/Neo*^ mice compared to WT. Plasma levels of hydrogen sulfide (H_2_S/HS^-^) tended to be higher in plasma from *Asl*^*Neo/Neo*^ mice versus WT (Supplementary Figure 1F) but did not differ in liver (Supplementary Figure 1G). In contrast, tissue total glutathione levels in the liver were significantly decreased (Figure 1L).

Sulfur-containing amino acids (and H_2_S) exist in different forms (29): in both, blood and tissues they occur in free form (either reduced as free thiol (R-SH) or oxidised as disulfide (R-SS-R), sulfenic (R-SOH), sulfinic (R-SO_2_H) or sulfonic acid (R-SO_3_H)) and bound (in the form of a mixed disulfide) to proteins. Sulfhydryl (SH) groups act as nanotransmitters and redox switches in cellular communication while also playing a protective role against oxidative stress by scavenging reactive oxygen species (29, 30). We therefore also measured free thiols in plasma and liver. In agreement with the total amounts of glutathione-related compounds described above, we observed significantly higher free homocysteine in plasma (Figure 1M) and liver (Figure 1N), homocystine in plasma (Supplementary Figure 1H), cysteine free thiols in plasma (Figure 1O) and liver (Figure 1P), cystine free thiols in plasma (Supplementary Figure 1I), and sulfide free thiols in plasma (Supplementary Figure 1J) and in liver (Supplementary Figure 1K). Homocystine and cystine levels in liver were below the limit of quantification of the assay. γ-glutamyl-cysteine was slightly higher in *Asl*^*Neo/Neo*^ mice versus WT in plasma (Supplementary Figure 1L) and in liver (Supplementary Figure 1M) while hepatic cysteinyl-glycine concentrations were higher in *Asl*^*Neo/Neo*^ compared to WT mice (Supplementary Figure 1N). The comparison of the different steady-state concentrations of precursors and breakdown products of glutathione suggested that precursors accumulate due to a bottleneck in one of the rate-limiting steps of glutathione biosynthesis while GSH catabolism by γ-glutamyltranspeptidase (y-GT) is enhanced, explaining the lower glutathione levels in both the circulation and liver of *Asl*^*Neo/Neo*^ mice.

To determine whether oxidative stress might further contribute to the lower levels of total glutathione in plasma and liver of *Asl*^*Neo/Neo*^ mice, we measured lipid peroxidation products using the thiobarbituric acid reactive substances (TBARS) assay in liver. This documented a moderate but significant increase in *Asl*^*Neo/Neo*^ mice versus WT (*p*=0.044, Figure 1Q). The steady-state concentration of the oxidative breakdown products of NO, nitrite (NO_2_^-^) and nitrate (NO_3_^-^), were significantly lower in plasma (Figure 1R) and liver (Figure 1S), as previously reported in this disorder (4, 5). Supporting NO deficiency, decreased nitroso-species were observed in *Asl*^*Neo/Neo*^ livers versus WT (Supplementary Figure 1O). The lack of difference in nitrotyrosine levels by western blot in *Asl*^*Neo/Neo*^ livers versus WT confirmed the absence of nitro-oxidative stress (Figure 1T, Supplementary Figure 1P). Altogether, these results indicate systemic glutathione depletion, contrasting with accumulation glutathione precursors, compromised NO production and moderate oxidative stress in ASA.

Glutathione biosynthesis relies on 2 enzymatic steps: (i) the rate limiting glutamate cysteine ligase (GCL) catalyses the conversion of L-glutamate + L-cysteine + ATP → γ-glutamyl-L-cysteine + ADP + Pi then (ii) glutathione synthase (GS) catalyses γ-glutamyl-L-cysteine + L-glycine ATP → glutathione + ADP + Pi. GCL is an heterodimer with a heavy catalytic subunit (GCLC) and a light or modifier subunit (GCLM) (19). Based on our findings, we hypothesised a deficiency in glutathione biosynthesis and confirmed downregulation of GCL with decreased gene expression of GCLC (Figure 1U) and GCLM (Figure 1V) but no significant decrease of GS (Figure 1W).

### *In vivo* evidence of impaired glutathione metabolism in *Asl*^*Neo/Neo*^ mice

The biosynthesis of glutathione is dependent on cellular import of cystine in exchange for glutamate efflux via the cystine/glutamate antiporter system X_C_^-^, a transmembrane transport system (Figure 2A). System X_C_^-^ is a heterodimeric protein, consisting of the major transporter xCT (*SLC7A11*) and CD98hc (*SLC3A2*) which is required for the membrane localization of multiple amino acid transporters. The radiolabelled glutamate analogue [^18^F]FSPG provides a functional readout of system X_C_-activity and corresponding *de novo* glutathione biosynthesis (31). Using positron emission tomography (PET), [^18^F]FSPG has been used in clinic for cancer diagnosis and preclinically to assess drug resistance in cancer (21, 22). In order to functionally assess alterations in the glutathione biosynthetic pathway, [^18^F]FSPG was administered intravenously (IV) to 2-3 weeks-old *Asl*^*Neo/Neo*^ mice and WT littermates, with radiotracer retention dynamically imaged by PET. Liver [^18^F]FSPG retention for *Asl*^*Neo/Neo*^ mice (14 ± 4% injected dose (ID)/g) was 3-fold higher (*p*=0.002) than that of WT mice (5.2 ± 1.5 %ID/g). In WT mice, the liver was barely visualized, with images being dominated by radiotracer retention in the pancreas and kidney, whereas it was challenging to distinguish between pancreatic and liver [^18^F]FSPG retention in *Asl*^*Neo/Neo*^ mice (Figures 2B, 2C, Supplementary Figure 2A). Unexpectedly, high [^18^F]FSPG retention was also present in the skin of *Asl*^*Neo/Neo*^ mice (13 ± 1.8 %ID/g) which was not the case in WT littermates (5.3 ± 2.3 %ID/g) (Figure 2F, Supplementary Figure 2B; *p*=0.002). We confirmed that the protein expression of xCT in 2 week-old *Asl*^*Neo/Neo*^ mice was substantially increased than that of the WT littermates, which had low baseline expression (Figures 2D). [^18^F]FSPG is therefore a useful non-invasive marker aberrant glutathione metabolism in the liver and skin of *Asl*^*Neo/Neo*^ mice, mediated at least in-part through the upregulated expression of the xCT antiporter.

**Figure 2.**
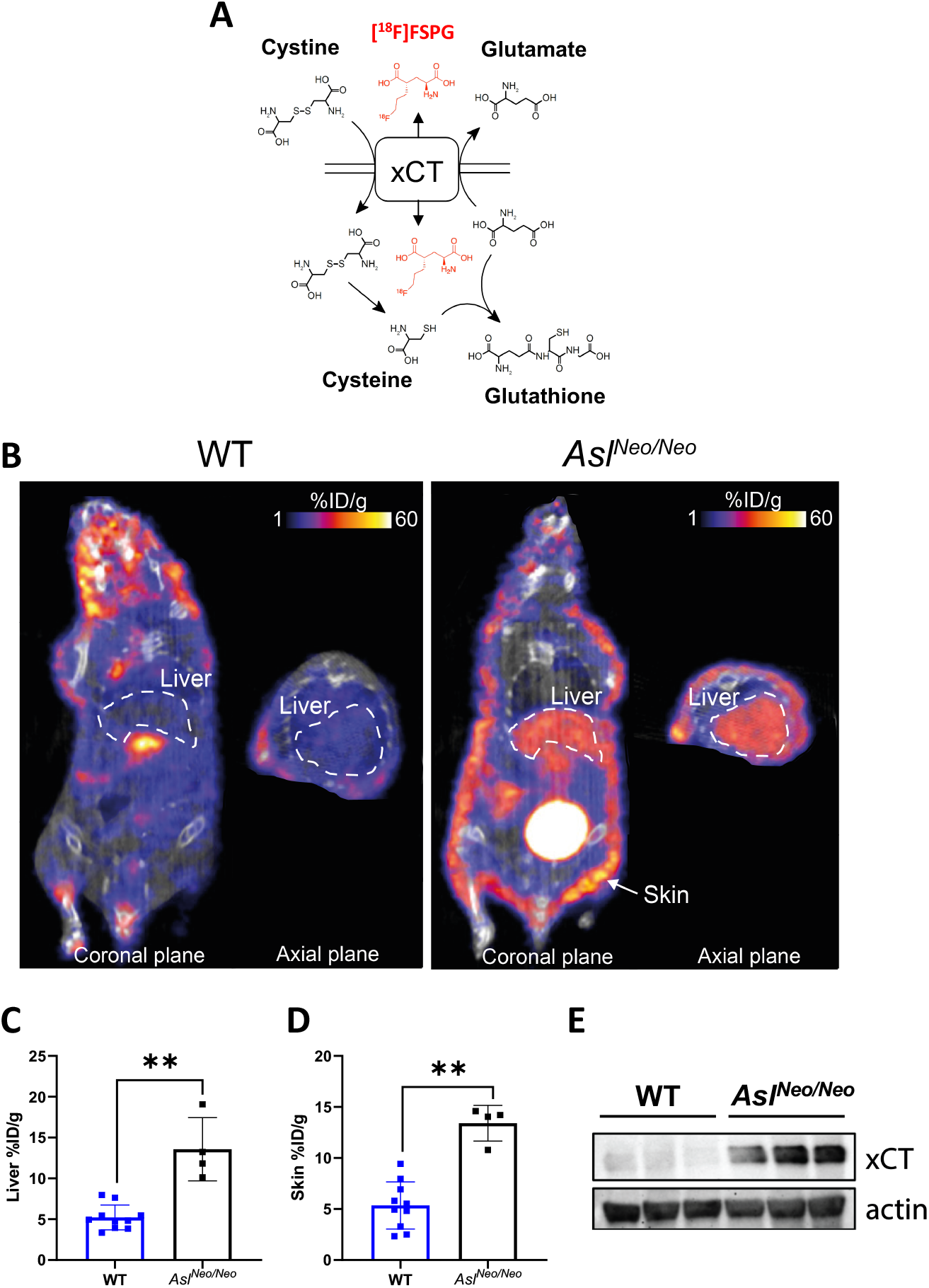
*In vivo* evidence of impaired glutathione metabolism in *Asl*^*Neo/Neo*^ mice. **(A)** Schematic overview of system X_C_^-^ function, shuttling cystine, glutamate and [^18^F]FSPG (red) across the cell membrane. Reduced cystine, cysteine, and glutamate are precursors for glutathione biosynthesis. **(B)** Representative PET/CT images of [^18^F]FSPG distribution (%ID/g), in the coronal and axial plane, of 2 week old WT littermates and *Asl*^*Neo/Neo*^ mice with increased [^18^F]FSPG retention present in the liver and skin of *Asl*^*Neo/Neo*^ mice. **(C, D)** Quantified [^18^F]FSPG retention in **(C)** the liver and **(D)** the skin of WT and *Asl*^*Neo/Neo*^ mice. **p<0.01, n=4-10. Graph shows mean ±SD. **(E)** Western blot of xCT expression, xCT is upregulated in the liver of *Asl*^*Neo/Neo*^ mice.

### Single intravenous administration of *hASL* mRNA corrects ureagenesis up to 7 days in *Asl*^*Neo/Neo*^ mice

The promising therapeutic effects of mRNA technology have been demonstrated recently in multiple liver inherited metabolic conditions (31, 32). Specifically engineered *hASL* mRNA encapsulated in lipid nanoparticles (LNP) were compared to *Luciferase (Luc)* mRNA used as control, which restored ASL expression and activity in ASL-deficient fibroblasts from patients compared to control after 24 and 48 hours (h) of incubation, respectively (Supplementary Figures 3A-C). mRNA therapy has transient efficacy and requires re-administration to enable sustained effect. Thus, a pharmacokinetic study of *hASL* mRNA *in vivo* was conducted in *Asl*^*Neo/Neo*^ mice to assess efficacy and duration of effect on the urea cycle. 3 week-old *Asl*^*Neo/Neo*^ mice received a single IV injection of either *hASL* or *Luc* mRNA at 1 mg mRNA/kg body weight and were sacrificed at 2 h, 24 h, 72 h or 7 days. A marked reduction of plasma ammonia (Figure 3A), argininosuccinic acid (Figure 3B) and citrulline (Figure 3C) in dried blood spots and urine orotate (Figure 3D) was observed within 24h of administration in *hASL* mRNA treated *Asl*^*Neo/Neo*^ mice compared to control (*Luc* mRNA) treated group. This effect was sustained over 7 days. Importantly, the levels of these metabolites were comparable in the *hASL* mRNA treated *Asl*^*Neo/Neo*^ mice to physiological levels in WT mice. Western blot and immunohistochemistry data in liver showed restored ASL protein expression at physiological levels at 24h post-administration of *hASL* mRNA (Figures 3E-H, Supplementary Figure 4A-C). ASL levels were consistently higher in *hASL* versus the *Luc* mRNA treated group. Liver ASL activity was also restored to physiological levels at 24h and 72h in *hASL* versus the *Luc* mRNA treated group, but began to decline by 7 days (Figure 3I).

**Figure 3.**
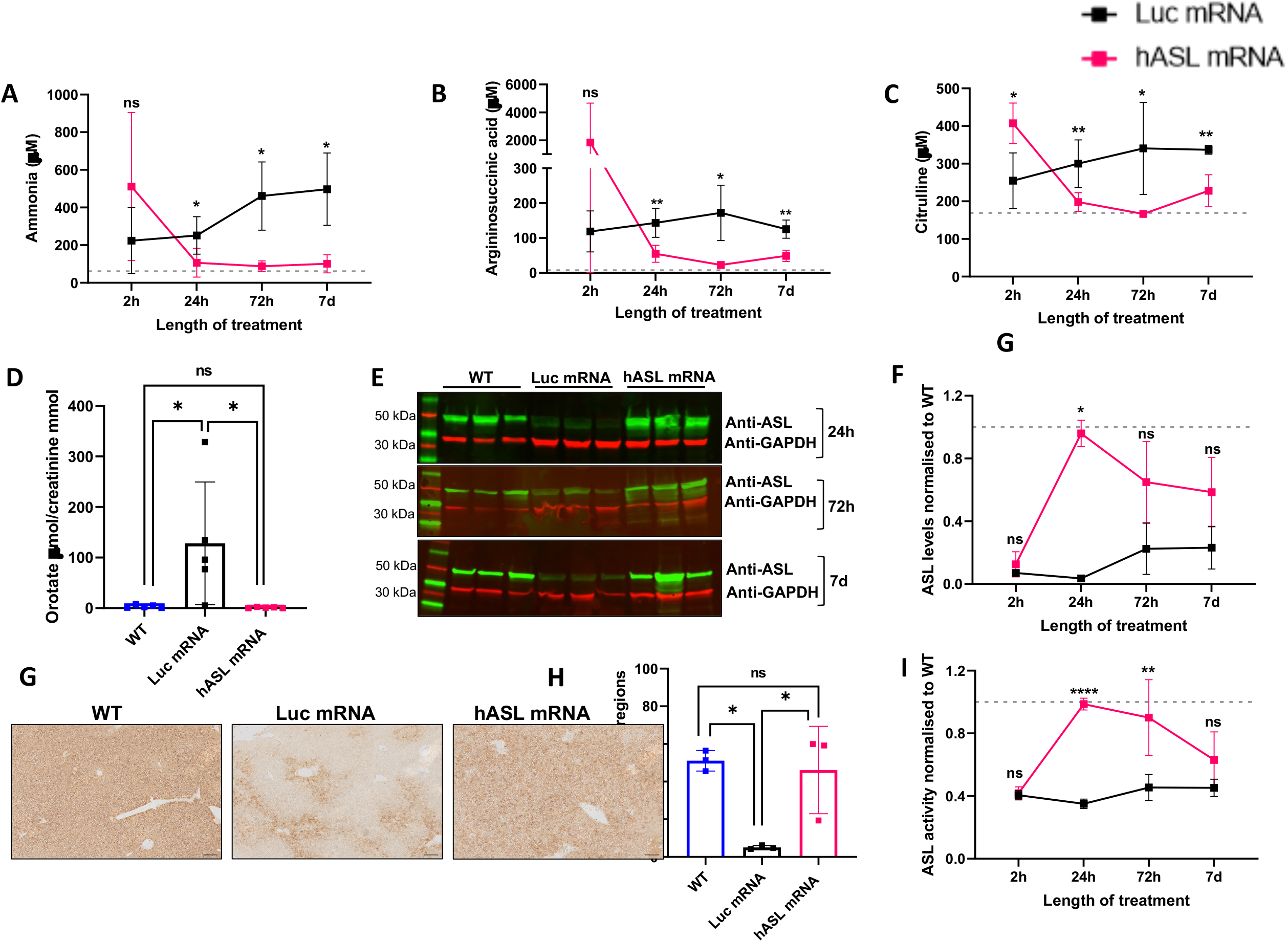
Pharmacodynamics of *hASL* mRNA *in vivo*. **(A**) Average ammonia levels from plasma and average (**B**) argininosuccinic acid (**C**) citrulline levels from dried blood spots at 2, 24, 72 hours and 7 days. (**D**) Urine orotic acid levels normalised to creatinine at 24 hours. (**E**) ASL western blot at 24 hours, 72 hours and 7 days. (**F**) Quantification of ASL immunoblot normalised to GAPDH (**G**) Representative images of liver ASL immunostaining at 24 hours post mRNA administration and (**H**) Quantification. Scale bar= 100μM. (**I**) Liver ASL activity at 2, 24, 72 hours and 7 days. Values normalised against WT control. (**A-D, F, H-I**) One-way ANOVA with Tukey’s post-test per timepoint, ns=not significant, *p<0.05, **p<0.01, ***p<0.005, ****p<0.0001. (**A-D, F, H-I**) n=3-7 per group. (**A-C, F, I**) Grey dotted line represents mean WT values. Graph shows mean +SD.

### *hASL* mRNA therapy from birth normalises the phenotype of *Asl*^*Neo/Neo*^ mice

Pharmacokinetic data showed a single mRNA dose to be efficacious up to 7 days. We initiated a study with repeated administration of *hASL* mRNA versus *Luc* mRNA in neonatal *Asl*^*Neo/Neo*^ pups. Mice received systemic administration of mRNA constructs every 7 days with the first IV dose administered at day 1 of life. Due to technical limitations, an intraperitoneal (IP) dose of 2 mg/kg at week 1 was performed based on equivalence of liver biodistribution between IV and IP routes, showing similar biodistribution with a two-fold higher IP dose compared to IV injection (Supplementary Figures 5A, 5B). Mice were treated for 7 weeks and harvested 48 h following the last injection (Figure 4A). The macroscopic phenotype of *Asl*^*Neo/Neo*^ mice was restored to that of WT littermates in the *hASL* mRNA treatment group with normalisation of survival (Figure 4B, *p*=0.002), growth (Figure 4C, Supplementary Figure 6A), fur (Figures 4D) and hepatomegaly (Supplementary Figure 6B). One *hASL* mRNA treated mutant was culled at 24 days of age due to malocclusion, a complication unrelated to the ASL phenotype or mRNA therapy. In contrast, *Luc* mRNA treated *Asl*^*Neo/Neo*^ littermates showed abnormal fur, impaired growth and early death within 2 weeks of life (Figures 4B-D).

**Figure 4.**
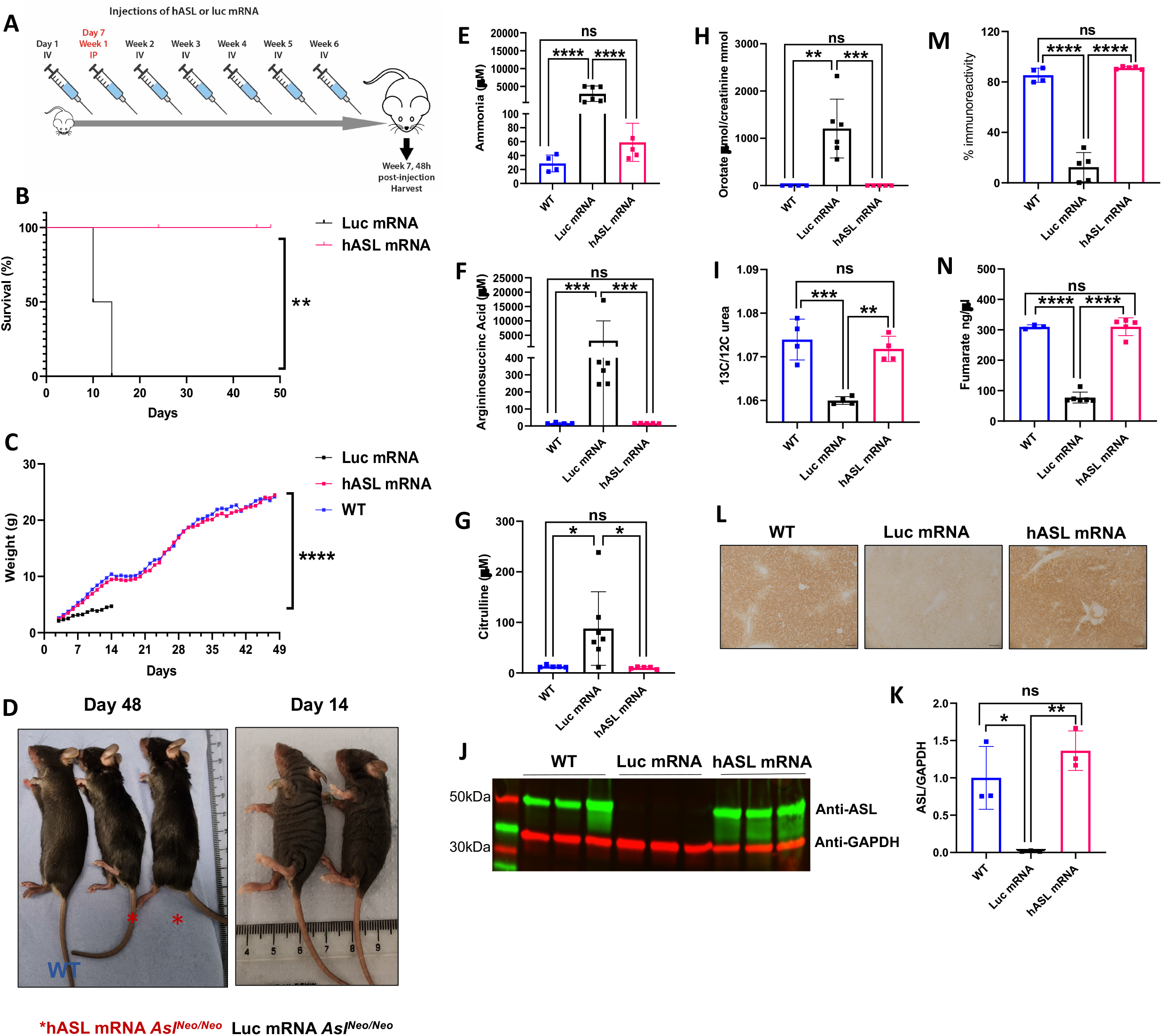
*hASL* mRNA therapy from birth corrects the phenotype of *Asl*^*Neo/Neo*^ mice. (**A**) Schematic illustration of experimental plan. *Asl*^*Neo/Neo*^ mice were given weekly intravenous (IV) dose of 1mg/kg of either *hASL* or *Luc* mRNA from birth up to 7 weeks, except for day 7 where the mice were administered intraperitoneally with dose of 2mg/kg. Harvest was performed 48 hours post the last injection. (**B**) Kaplan-Meier survival curve of *hASL* and *Luc*-mRNA treated *Asl*^*Neo/Neo*^ mice. (**C**) Average growth curve of WT, *hASL* and *Luc*-mRNA treated *Asl*^*Neo/Neo*^ mice. (**D**) Representative images of WT, *hASL* and *Luc* mRNA treated *Asl*^*Neo/Neo*^ mice at harvest. (**E**) Average plasma ammonia concentration, (**F**) argininosuccinic acid (**G**) and citrulline concentrations from dried blood spots, (**H**) urine orotic acid and (**I**) C13 ureagenesis from WT, *hASL* and *Luc*-mRNA treated *Asl*^*Neo/Neo*^ mice. (**J**) ASL western blot of WT, *hASL* and *Luc*-mRNA treated *Asl*^*Neo/Neo*^ mice and (**K**) quantification. (**L**) Representative images of ASL immunostaining in livers of WT, *hASL* and *Luc*-mRNA treated *Asl*^*Neo/Neo*^ mice and (**M**) quantification. (**N**) Liver ASL activity from WT, *hASL* and *Luc*-mRNA treated *Asl*^*Neo/Neo*^ mice livers (**D**) Scale bar=2cm. (**L**) Scale bar= 100μM. (**B**) Log-rank (Mantel-Cox) (**C**) One-way ANOVA with Sidak’s post-test comparison. (**E-I, K, M, N**) One-way ANOVA with Tukey’s post-test analysis, ns=not significant, *p<0.05, **p<0.01, ***p<0.005, ****p<0.0001. **(C-H, M, N)** WT n=4; *hASL* mRNA n=5; *Luc* mRNA n=5-6. Graph shows mean +SD.

Animals which survived 7 weeks were culled 48 h after the last mRNA injection. Analysis showed normalization of ammonaemia (Figure 4E), argininosuccinic acid (Figure 4F) and citrulline (Figure 4G) levels in dried blood spots, and urinary orotate levels (Figure 4H) in *hASL* mRNA treated *Asl*^*Neo/Neo*^ mice. Elevated plasma amino transferase (ALT) was normalized in the *hASL* mRNA treated *Asl*^*Neo/Neo*^ mice (Supplementary Figure 6C). Longitudinal analysis of plasma ammonia levels (Supplementary Figure 6D), argininosuccinic acid (Supplementary Figure 6E) and citrulline (Supplementary Figure 6F) levels in dried blood spots, and urinary orotate levels (Supplementary Figure 6G,I) in *hASL* mRNA treated *Asl*^*Neo/Neo*^ mice showed sustained therapeutic benefit over time. Next, functional assessment of urea cycle *in vivo* was measured by quantifying labelled urea in the plasma following the injection of ^13^C labelled sodium acetate 30 minutes pre-harvest. ^13^C labelling showed restored ureagenesis in the *hASL* treated *Asl*^*Neo/Neo*^ mice (Figure 4I). ASL levels in liver assessed by western blot (Figures 4J, 4K) and immunohistochemistry (Figures 4L, 4M) were restored to physiological levels and physiological pattern following *hASL* mRNA therapy. Importantly, ASL activity in liver was restored to WT physiological levels following *hASL* mRNA therapy in *Asl*^*Neo/Neo*^ mice (Fig 4N).

### *hASL* mRNA therapy partially rescues the adult phenotype in *Asl*^*Neo/Neo*^ mice

Next, we assessed the rescue of the severe phenotype of *Asl*^*Neo/Neo*^ mice following late initiation of *hASL* mRNA therapy. *Asl*^*Neo/Neo*^ mice received their first IV *hASL* mRNA dose in early adulthood at day 21 of life followed by weekly mRNA administration for up to 9 weeks (Figure 5A). All treated mice survived to the end of the study except one that died after 2 injections at day 31 of life, while *Luc* mRNA treated mice only survived up to day 37 of life with most animals dying before day 30 of life (Figure 5B, *p*=0.0025). *hASL* mRNA treated *Asl*^*Neo/Neo*^ mice showed significantly improved growth compared to *Luc* mRNA *Asl*^*Neo/Neo*^ littermates, however the body weight remained significantly lower than WT littermates (Figure 5C Supplementary Figure 7A). The full recovery of hair growth in *hASL* mRNA treated *Asl*^*Neo/Neo*^ mice was observed with similar fur pattern compared to WT (Figures 5D). Liver-to-body weight ratio remained significantly elevated in both *Asl*^*Neo/Neo*^ mice groups (Supplementary Figure 7B). Plasma ammonia (Figure 5E), argininosuccinic acid (Figure 5F) and citrulline (Figure 5G) in dried blood spots and urinary orotic acid levels (Figure 5H) were significantly reduced following *hASL* mRNA therapy to physiological WT levels. Importantly, ^13^C ureagenesis analysis showed restored *in vivo* urea cycle function compared to that of WT physiological levels in *hASL* mRNA treated *Asl*^*Neo/Neo*^ mice (Figure 5I) compared to *Luc* mRNA treated *Asl*^*Neo/Neo*^ littermates. ALT analysis indicated an absence of liver toxicity in the *hASL* mRNA treated *Asl*^*Neo/Neo*^ mice compared to the WT littermates (Supplementary Figure 7C). ASL levels in liver assessed by western blot (Figure 5J, K) were restored to physiological levels following *hASL* mRNA therapy compared to WT. Liver ASL activity was significantly improved following *hASL* mRNA compared to *Luc* mRNA in *Asl*^*Neo/Neo*^ mice (Figure 5L).

**Figure 5.**
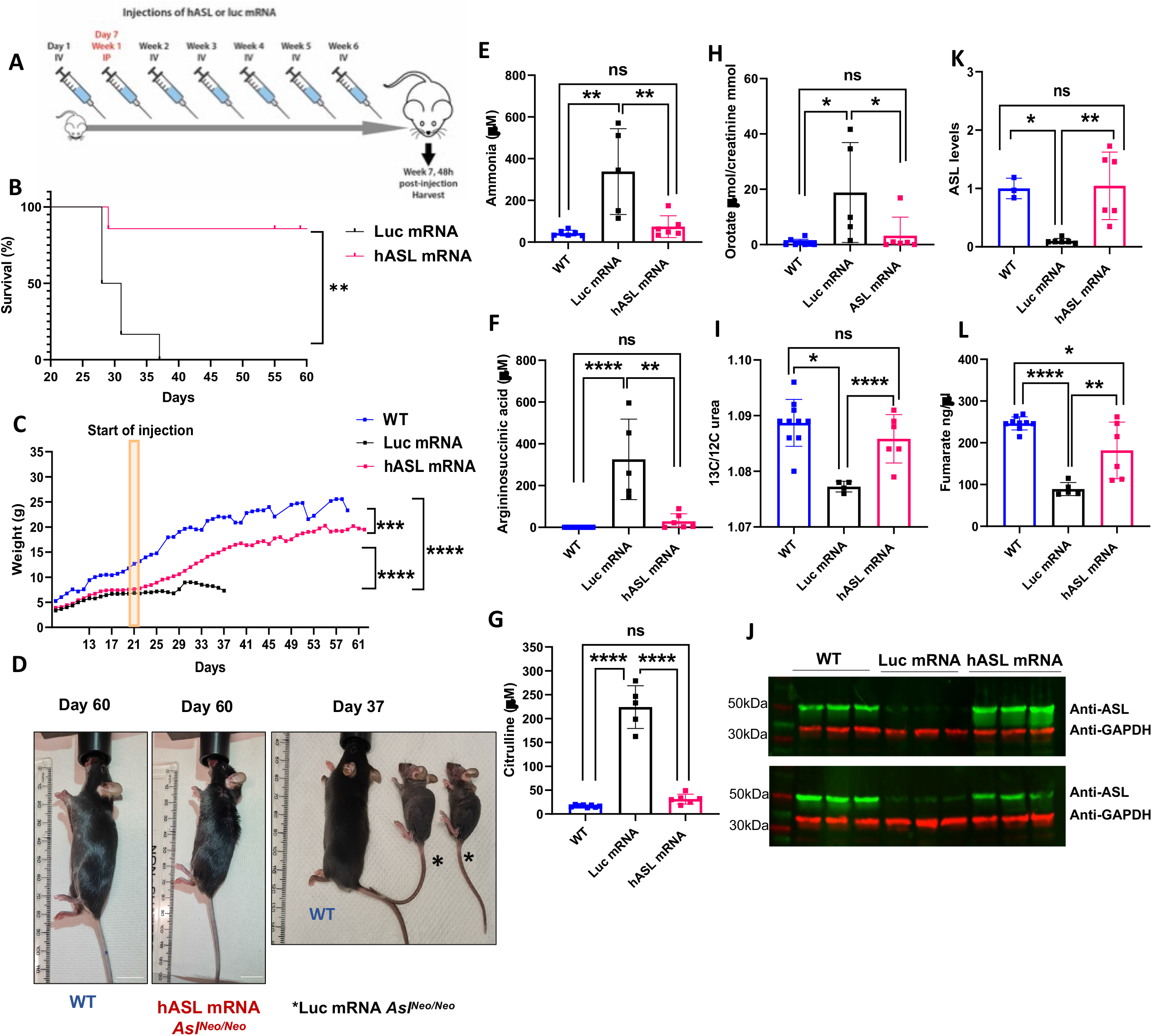
*hASL* mRNA therapy partially rescues the adult phenotype in *Asl*^*Neo/Neo*^ mice. (**A**) Schematic illustration of experimental plan. *Asl*^*Neo/Neo*^ mice were given weekly intravenous (IV) dose of 1mg/kg of either *hASL* or *Luc* mRNA from day 21 up to 9 weeks. (**B**) Kaplan-Meier survival curve of *hASL* and *Luc*-mRNA treated *Asl*^*Neo/Neo*^ mice. (**C**) Average growth curve of WT, *hASL* and *Luc*-mRNA treated *Asl*^*Neo/Neo*^ mice. (**D**) Representative images of WT, *hASL* and *Luc* mRNA treated *Asl*^*Neo/Neo*^ mice at harvest. (**E**) Average plasma ammonia concentration, (**F**) argininosuccinic acid (**G**) and citrulline concentrations from dried blood spots, (**H**) urine orotic acid and (**I**) C13 ureagenesis from WT, *hASL* and *Luc*-mRNA treated *Asl*^*Neo/Neo*^ mice. (**J**) ASL western blot of WT, *hASL* and *Luc*-mRNA treated *Asl*^*Neo/Neo*^ mice and (**K**) quantification (**L**) Liver ASL activity from WT, *hASL* and *Luc*-mRNA treated *Asl*^*Neo/Neo*^ mice livers (**B**) Log-rank (Mantel-Cox), *p*=0.0025 (**C**) One-way ANOVA with Sidak’s post-test comparison. (**D**) Scale bar=2cm. (**E-I, K, L**) One-way ANOVA with Tukey’s post-test analysis, ns=not significant, *p<0.05, **p<0.01, ****p<0.0001. **(C-D, E-H, K, L)** WT n=6, *Luc* mRNA n=4-6, *hASL* mRNA n=6-7 Graph shows mean +SD.

### *hASL* mRNA therapy corrects the metabolic dysfunction and liver pathophysiology in *Asl*^*Neo/Neo*^ mice

Next, we wanted to determine the extent of the correction of liver metabolic dysfunction following *hASL* mRNA treatment in *Asl*^*Neo/Neo*^ mice using RNA-sequencing (RNA-seq) transcriptomic analysis. We began by visualising the overall variation in gene expression across WT and *Asl*^*Neo/Neo*^ mice treated at birth with either *hASL* mRNA or *Luc* mRNA therapy. Principal component analysis showed clustering of the *hASL* mRNA treated and WT liver samples, suggesting a similar profile of gene expression in both groups. In contrast, *Luc* mRNA treated *Asl*^*Neo/Neo*^ mice clustered separately (Figure 6A). Next, we analysed differential gene expression identifying all genes with a log2-fold change of >0.1 or < -0.1 and passing an FDR cut off of <0.05. Comparing WT vs *Luc* mRNA *Asl*^*Neo/Neo*^ groups, we found 2705 genes to be significantly up- (1257 genes) or down-regulated (1448 genes) (Figure 6B). Remarkably, only 7 genes (1 upregulated and 6 downregulated) were differentially expressed between WT vs *hASL* mRNA *Asl*^*Neo/Neo*^ mice livers, thereby demonstrating the significant efficacy of mRNA therapy in correcting liver dysfunction (Figure 6C). This interpretation of the data was supported by the analysis of differential gene expression between *Luc* mRNA and *hASL* mRNA *Asl*^*Neo/Neo*^. Similar to the WT vs *Luc* mRNA *Asl*^*Neo/Neo*^ comparison (Figure 6D), we identified a large number of differentially expressed genes (4297 genes) with 1962 genes being significantly upregulated (log2-fold change > 0.1 and FDR < 0.05) and 2335 being significantly downregulated (log2-fold change < -0.1 and FDR < 0.05). Of note, the murine *Asl* gene was significantly downregulated in both groups of *Asl*^*Neo/Neo*^ livers, likely due to the codon optimised mRNA sequence being distinct from that contained within the mouse transcriptome build used for sequence alignment.

**Figure 6.**
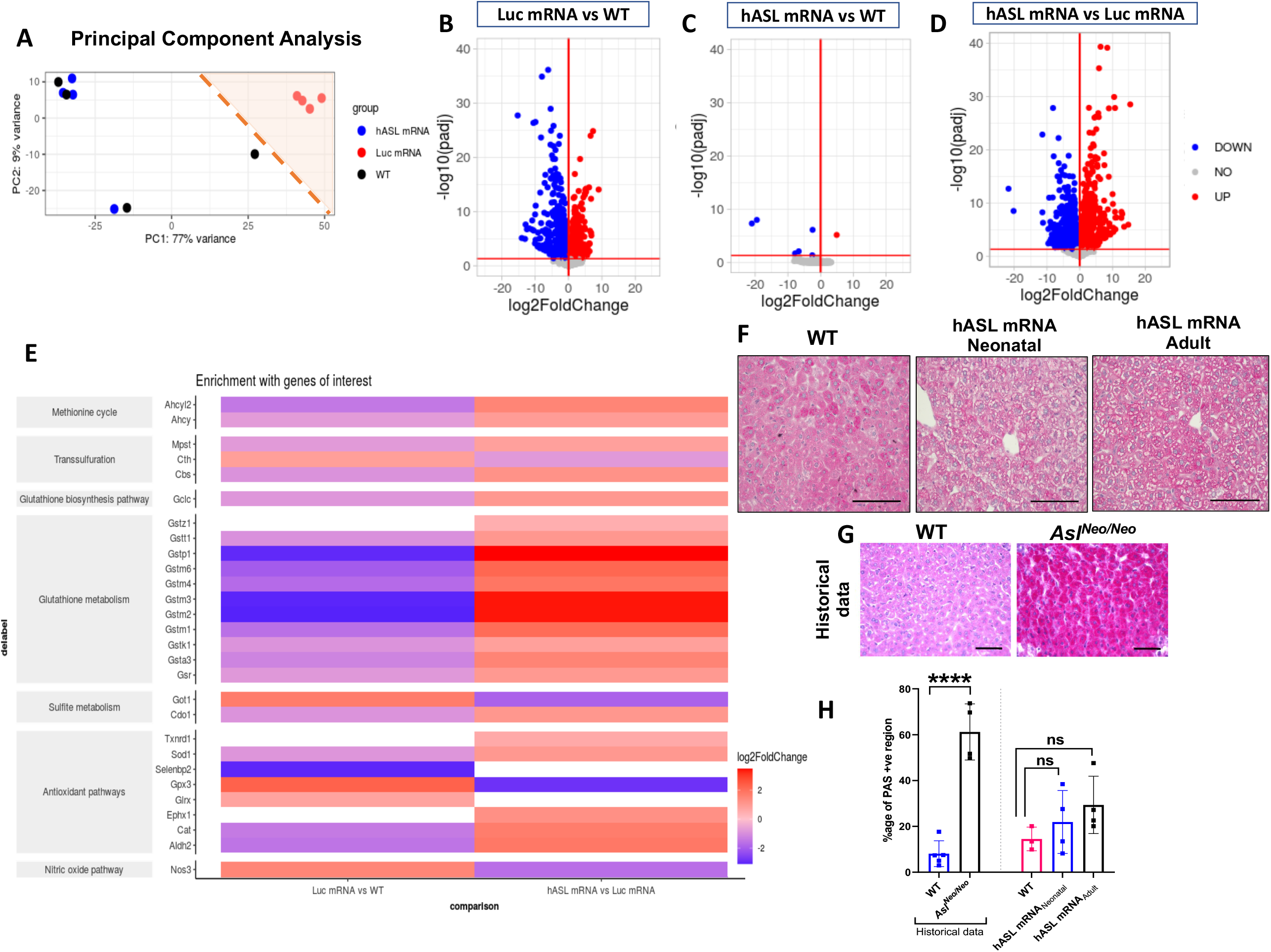
*hASL* mRNA therapy corrects the metabolic dysfunction and liver pathophysiology in *Asl*^*Neo/Neo*^ mice. (**A**) Principal component analysis plots comparing treatment applied (untreated WT, *hASL* or *Luc* mRNA) and mouse genotype (WT or *Asl*^*Neo/Neo*^) with percentage of variance associated with each axis. (**B**) Volcano plots showing differential gene expression (DEG) analysis of *Luc* mRNA vs WT, (**C**) *hASL* mRNA vs WT and (**D**) *hASL* mRNA vs *Luc* mRNA. Scatter plots show log transformed adjusted p-values (<0.05) on the y-axis against log2 fold change (>0.10) values on the x-axis. Blue and red dots represent genes that are significantly downregulated and upregulated respectively between groups. Grey dots represent genes that are not significantly altered. (**E**) Pathway analysis highlighting genes of interest significantly altered in DEG analysis organised with their associated pathways when comparing *Luc* mRNA vs WT and *hASL* mRNA vs *Luc* mRNA groups. (**F**) Periodic Acid Schiff staining of liver samples harvested from WT and *hASL mRNA* treated *Asl*^*Neo/Neo*^ mice from neonatal or adulthood. (**G**) Historical Periodic Acid Schiff staining of liver samples harvested from WT and *Asl*^*Neo/Neo*^ mice treated with vehicle. (**F**) Scale bar: 100 μm. (**G**) Scale bar: 500 μm. **(H)** Quantification of hepatic glycogen from immunostaining. (**F, H**) One way ANOVA with Tukey’s post-test, ns=not significant, (**G, H**) Unpaired 2-tailed Student’s t test, ****p<0.0001. (**F, H**) WT n=3, *hASL mRNA* treated *Asl*^*Neo/Neo*^ mice n=4 (**G, H**) WT n= 5, *Asl*^*Neo/Neo*^ n= 4. Graph shows mean +SD.

To further study the dysregulation of glutathione function, the analysis of pathways affecting glutathione metabolism was performed on the RNA-seq data. Our analysis highlighted downregulation of multiple genes involved in glutathione biosynthesis and metabolism alongside alterations of genes of the methionine cycle, transsulfuration and antioxidant pathways between *Luc* mRNA (control) *Asl*^*Neo/Neo*^ mice and WT livers (Figure 6E). These findings additionally support disruption of glutathione metabolism in *Asl*^*Neo/Neo*^ mice. Importantly, these pathways were corrected post *hASL* mRNA treatment as shown by post pathways analysis comparison between *hASL* mRNA and *Luc* mRNA treated *Asl*^*Neo/Neo*^ mice (Figure 6E).

*Asl*^*Neo/Neo*^ mice recapitulate chronic ASA liver pathology with progressive glycogen deposition compared to WT (8, 11, 16). To assess whether prolonged mRNA therapy could correct the chronic liver disease in ASA, we compared glycogen storage in the livers of age-matched adult WT and *hASL* treated mice using Periodic Acid Schiff (PAS) staining. As *Luc* mRNA treated *Asl*^*Neo/Neo*^ mice died early before adulthood, their liver did not present with glycogen deposition at harvest. The data was then compared against historical data from *Asl*^*Neo/Neo*^ mice which survived longer owing to supportive treatment of sodium benzoate and arginine supplementation (Figures 6F-H). Historical data from *Asl*^*Neo/Neo*^ mice showed 7.5-fold higher glycogen accumulation compared to their age-matched WT littermates (*p*<0.0001) (*Asl*^*Neo/Neo*^ mice = 61.22 ± 12.24%; WT = 9.08 ± 5.6%) (11). In our current comparison, *Asl*^*Neo/Neo*^ mice treated from birth with *hASL* mRNA showed no significant difference in glycogen accumulation compared to their WT littermates (WT = 4.51 ± 5.2%, *hASL* mRNA neonatal = 21.91 ± 13.7%, *p*=0.69). Similarly, *Asl*^*Neo/Neo*^ mice treated at adulthood showed no significant difference (29.37 ± 12.5%, *p*=0.69). Thus, mRNA therapy improves glycogen deposition in ASA, contributing to the improvement of chronic hepatocellular injury.

### *hASL* mRNA therapy corrects the dysfunction of glutathione metabolism in *Asl*^*Neo/Neo*^ mice

To investigate the potential of [^18^F]FSPG PET as a non-invasive tool in assessing therapeutic efficacy, [^18^F]FSPG was administered IV to 2 weeks-old untreated and *hASL* mRNA treated *Asl*^*Neo/Neo*^ livers (IV administration of 1 mg/kg *hASL* mRNA at birth followed by weekly IP administration of 2 mg/kg mRNA before imaging at 2 weeks of age). Supporting our functional and metabolic data, [^18^F]FSPG retention was halved in *hASL* mRNA treated (11 ± 2.0 %ID/g) versus untreated *Asl*^*Neo/Neo*^ mice (22 ± 2.3 % ID/g; *p* = 0.026) (Figures 7A, 7B; Supplementary Figure 8A), corresponding with a restoration of liver glutathione for both neonatal and adult treated *Asl*^*Neo/Neo*^ mice (Figure 7C). However, [^18^F]FSPG retention was not completely restored to baseline levels from WT livers (5.0 ± 2.8 %ID/g). This restoration of glutathione levels was associated with a significant reduction of total homocysteine ratio compared to WT in liver from *hASL* mRNA treated versus *Asl*^*Neo/Neo*^ mice (Figure 7D). In line with improvement of glutathione metabolism and [^18^F]FSPG retention, the expression of cystine/glutamate antiporter system X_C_^-^ was massively reduced in livers from *hASL* mRNA treated versus *Asl*^*Neo/Neo*^ mice (Figure 7E).

**Figure 7.**
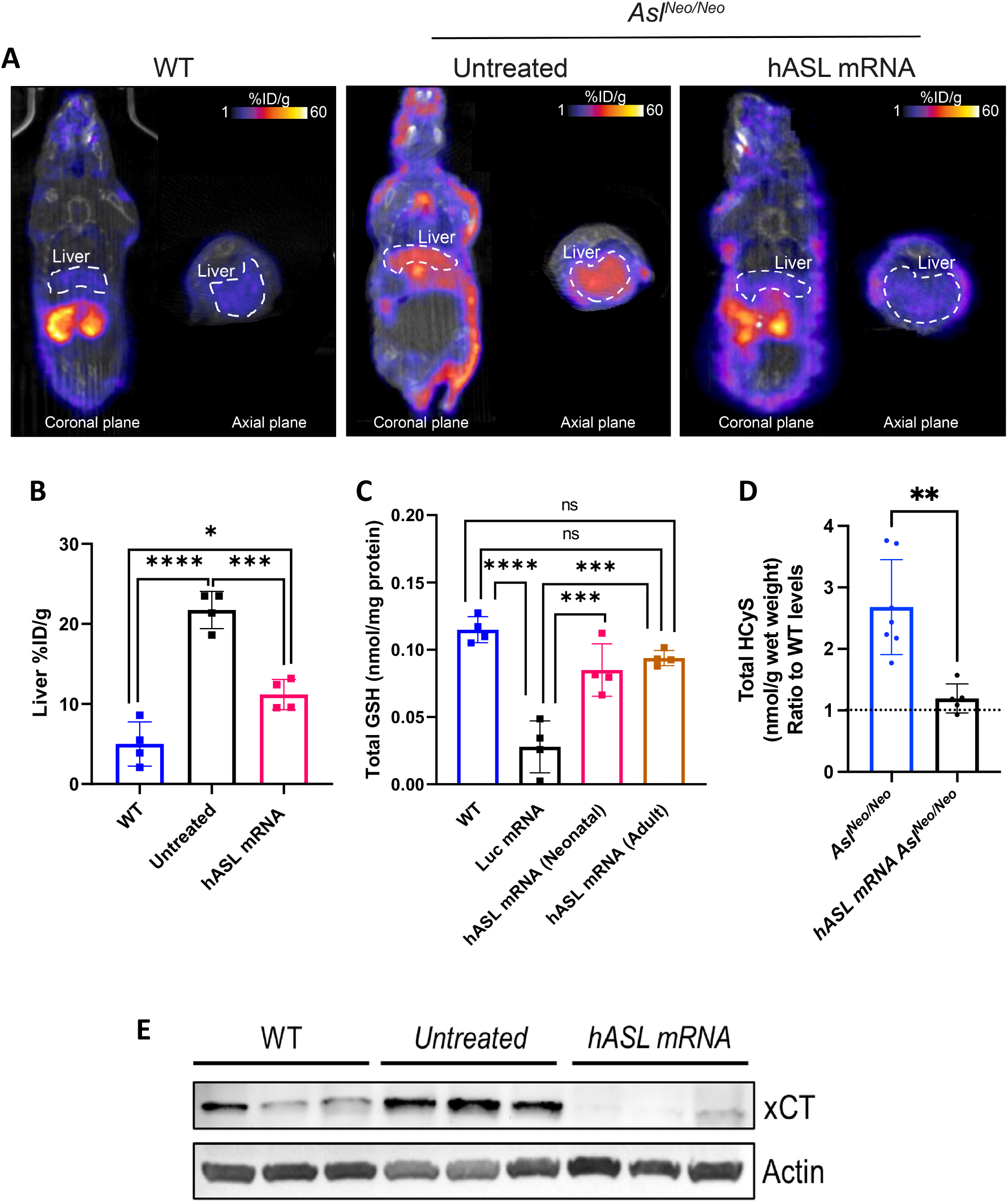
*hASL* mRNA therapy corrects the dysfunction of glutathione metabolism in *Asl*^*Neo/Neo*^ mice. **(A)** [^18^F]FSPG distribution (%ID/g) in representative coronal and axial plane PET/CT images of WT, untreated *Asl*^*Neo/Neo*^ and *hASL* mRNA treated *Asl*^*Neo/Neo*^ mice. **(B)** [^18^F]FSPG quantification of the liver in WT, untreated *Asl*^*Neo/Neo*^ and *hASL* mRNA treated *Asl*^*Neo/Neo*^ mice. **(C**) Total glutathione levels from liver in WT, Luc mRNA treated *Asl*^*Neo/Neo*^ mice and *hASL* mRNA treated *Asl*^*Neo/Neo*^ mice from neonatal or adulthood. **(D)** Liver total homocysteine concentrations expressed as ratio out of WT levels (shown as dotted line) from untreated versus *hASL* mRNA treated *Asl*^*Neo/Neo*^ mice in adulthood. **(E)** Western blot of xCT expression, upregulated xCT in untreated *Asl*^*Neo/Neo*^ liver is decreased in the liver of *hASL* mRNA treated *Asl*^*Neo/Neo*^ mice. Graph shows mean ±SD. One-way ANOVA with Tukey’s post-test, ns=not significant, ***p<0.001, ****p<0.0001, n=4-6.

Conversely, skin [^18^F]FSPG retention was not affected by mRNA therapy. [^18^F]FSPG skin retention was 4.2 ± 3.4 %ID/g in WT mice, 15 ± 3.7 %ID/g and 15 ± 4.2 %ID/g in untreated and *hASL* mRNA treated *Asl*^*Neo/Neo*^ mice, respectively (Supplementary Figures 8B, 8C).

## Discussion

A chronic liver involvement has been observed in most UCDs, with hepatomegaly, elevated transaminases, chronic liver dysfunction, steatosis and/or glycogen accumulation (7, 33). Among all UCDs, ASL deficiency is reported with a greater burden as well as higher frequency and severity of liver complications (1, 8, 9, 33). Liver symptoms are commonly observed in 37-49% of ASA patients with a higher frequency in early-onset versus late-onset patients. Liver symptoms commonly include hepatomegaly and transaminitis, but can show chronic liver failure, glycogen deposition, fibrosis, and hepatocellular carcinoma (6-10, 12, 13). The *Asl*^*Neo/Neo*^ mouse model recapitulates much of the human phenotype of chronic hepatocellular injury with reports of hepatomegaly, elevated transaminases, aberrant hepatic glycogen accumulation and variable fibrosis (8, 11, 16, 18). The liver pathology persists and can worsen despite appropriate ammonia control under standard of care, suggesting hyperammonaemia is not the sole cause (7). Accumulation of toxic metabolites such as argininosuccinic acid, potentially facilitated by high arginine supplementation could be contributing factors (6, 9, 34).

Oxidative stress has been described in UCDs, mostly in the context of acute hyperammonaemia triggering calcium signalling, activating N-methyl-D aspartate (NMDA) receptors, activating NOS and promoting the production of reactive nitrosative and oxidative species in the brain (35). *In vitro* studies have suggested the pro-oxidative role of amino acid imbalance, such as excess of ornithine, citrulline or arginine in different UCDs (36). Oxidative stress is a well-recognised pathophysiological mechanism observed in ASA. Increased oxidative stress either systemically (5), affecting neuronal (16) or endothelial (28) cells has been described, potentially caused by either low arginine and NOS uncoupling (16, 34) or direct toxicity from argininosuccinic acid and derived conjugates like guanidinosuccinate (14, 34, 37). Additionally, ASL deficiency is a recognised model of inherited endothelial dysfunction due to compromised NO production (28). Defective NO-related vasorelaxation causes arterial hypertension which can be restored by nitrite supplementation (5). Endothelial dysfunction associated with oxidative stress is likely to play a role in the pathophysiology of the liver disease. However, in our study the *Asl*^*Neo/Neo*^ mouse presents only moderate evidence of hepatic oxidative stress with mildly raised lipid peroxidation, but without increased nitrotyrosine levels and lower systemic NO production as evidenced by reduced concentrations of nitrate and nitrite. Since NO is also a chain-breaking antioxidant capable of attenuating lipid peroxidation (38). ASL deficiency would seem to result in a double burden of enhanced oxidant stress as a result of both glutathione depletion and NO deficiency.

Since glutathione is a key antioxidant, mitochondrial protectant and master regulator of multiple redox processes (39, 40), its intracellular concentrations are tightly regulated. While acute oxidative stress typically leads to a transient shift in the GSH/GSSG ratio, sustained oxidative stress during chronic liver diseases such as cholestasis and toxic liver disease is often associated with compromised glutathione biosynthesis and/or a depletion of total tissue glutathione levels (19). We observed lower overall glutathione levels due to a combination of compromised biosynthesis and loss due to increased breakdown with limited evidence for marked oxidative stress in *Asl*^*Neo/Neo*^ livers. These changes were accompanied by the upregulation of intracellular cystine uptake via the xCT antiporter, the increase of total and free thiol content of upstream metabolites of glutathione biosynthesis and the downregulation of genes involved in antioxidant pathways. Clearly, glutathione depletion in ASA cannot be explained by either deficiency of upstream precursors such as cysteine or severe oxidative stress.

Glutathione biosynthesis is an energy-dependent two-step enzymatic process whereby GCL catalyses the conversion of L-glutamate and L-cysteine to produce the c-glutamyl-L-cysteine, which is subsequently converted to glutathione by addition of L-glycine by GS (19). GCL activity is the rate limiting step in glutathione biosynthesis and requires GCLC (catalytic subunit) and GCLM (functional modulator subunit) (19, 41). In our work, the accumulation of γ-glutamylcysteine was not as marked as other glutathione precursors such as homocysteine and cysteine, which is consistent with the downregulation of GCLC in the transcriptomic dataset, and the decreased expression of both GCLC and GS by qRT-PCR. As glutathione synthesis is ATP-dependent (42), energy depletion and mitochondrial dysfunction, previously reported in ASA (28, 43), may further contribute to overall reduced glutathione biosynthesis. Decrease in Mg^2+^ or Mn^2+^ availability, required for the first catalytic step of glutathione synthesis could also play a role (41), as well as dysregulation of the transcriptional factor nuclear factor-erythroid 2 p45-related factor 2 (Nrf2), known to regulate expression of GCL and other antioxidant pathways (44). A multifactorial combination could therefore account for the severity of liver glutathione depletion with both decrease of glutathione biosynthesis and accelerated loss.

Glutathione as an essential antioxidant has multiple functions in scavenging free radicals, detoxification of electrophiles and maintenance of protein thiol status thereby regulating cellular survival, signalling and metabolism (19, 45). Glutathione depletion has been associated with the progression of liver diseases (45). Additionally, hyperhomocysteinemia has been reported with impaired liver function including liver failure, steatosis or fibrosis in patients (46, 47) and in animal models of homocystinuria (48, 49). While the exact mechanism remains unknown, the triad of glutathione depletion, hyperhomocysteinaemia and occasional hyperammonaemic events might explain the chronic liver phenotype in ASA, although further studies will be required to unravel the pathophysiological role of each of these mechanisms.

The upregulation of the xCT transporter in the liver suggests a feedback mechanism to alleviate the consequences of glutathione depletion. This mechanism is adopted by cancer cells to promote glutathione biosynthesis and thereby cell survival from an increase of the intracellular cysteine pool (50, 51). Large increases in xCT in *Asl*^*Neo/Neo*^ mice indicate that the liver shares the same plasticity. An unexpected finding was the increase in the xCT transporter function in the skin of the *Asl*^*Neo/Neo*^ mice, which may explain the skin inflammation observed in some patients and the hair and fur abnormalities noted in the *Asl*^*Neo/Neo*^ mouse, highlighting the multiorgan pathophysiology of ASA (52, 53).

This study highlights the potential of non-invasive [^18^F]FSPG-PET imaging as a sensitive diagnostic tool to assess the liver disease and investigate its pathophysiology in ASA. [^18^F]FSPG-PET imaging has been used in cancer clinical setting for diagnostic purposes and in preclinical studies to predict response to chemotherapy (21, 22). While the use of PET tracers in monogenic diseases is an unusual application in this disorder and in the context of active development of liver-targeting gene therapies, it could nevertheless become a valuable companion diagnostic to assess effective correction of the impaired glutathione metabolism in ASA liver. Elevated radiotracer retention in the skin further highlights the advantages of using whole-body imaging to reveal new insights into this, and other pathologies that disrupt glutathione homeostasis.

The lack of effective therapies for both ureagenesis and the chronic liver disease in ASA has promoted the development of various novel experimental therapies. The autophagy enhancer Tat-Beclin-1 (TB-1) peptide has shown partial increased survival and improved ureagenesis along with reduction in hepatocellular injury and glycogen accumulation that may prevent long-term hepatotoxicity (11). Restoration of ureagenesis was achieved using liver-targeting viral-mediated gene therapies using adenoviral (5, 8), adeno-associated viral (AAV) (16, 18), or lentiviral vectors (17). For viral vector-mediated gene delivery, there are ongoing concerns over capsid immunogenicity and toxicity. There have been recent reports of serious hepatotoxicity following AAV gene delivery, in XLMTM & SMN (54-57). In parallel, lipid nanoparticles encapsulating mRNA is emerging as a promising therapy for liver monogenic diseases (58-60). Through encapsulation of mRNA in optimised LNPs, functional therapeutic protein can be delivered to target cells with comparatively longer half-life and lower costs than protein replacement therapies. The absence of LNP immunogenicity and integration in the host genome enables safe repeating administration to compensate for mRNA degradation and transient efficacy and is becoming a safe alternative to viral vectors (61, 62). Proof of concept in liver inherited metabolic diseases has been rapidly increasing (31, 32, 63, 64) with supporting data for early phase clinical trials for ornithine transcarbamylase (NCT04442347), propionic acidaemia (NCT04899310) and methylmalonic acidaemia (NCT04159103).

Based on previous proof of concept studies performed in arginase deficiency, another UCD (65, 66), the therapeutic dose of 1 mg/kg of *hASL* mRNA delivered weekly through systemic routes of administration was tested in the *Asl*^*Neo/Neo*^ mouse. The treatment of animals from birth showed remarkable effects by normalising the macroscopic phenotype, biomarkers, *in vivo* ureagenesis, liver ASL expression and activity to physiological levels. Comparatively, induction of therapy in early adulthood showed partial but still very effective phenotypic rescue. Overall, this study showed proof of concept of mRNA therapy for both early-onset or rescue therapy in ASA and is actively supporting clinical translation.

Interestingly, dysregulation of glutathione metabolism and antioxidant pathways were fully corrected following mRNA therapy as observed in liver transcriptomics and liver glutathione levels (11). Similarly liver glycogen metabolism was well corrected. The lack of hepatomegaly in the neonatally-treated group and no increase in the plasma transaminase alanine amino transferase (ALT) in both treated groups also supported the correction of long-term liver pathology. The liver [^18^F]FSPG retention in neonatally-treated animals was halved compared to untreated mice, however not completely normalised. This could be due to the experimental design, which included a short 2-weeks period of therapy to achieve age-matched comparison with untreated control *Asl*^*Neo/Neo*^ animals. Interestingly, the correction was limited to liver with no benefit observed in the skin, demonstrating the liver targeting of mRNA therapy.

In conclusion, our study provides new insight into the liver pathophysiology of ASA, a rare metabolic disease, and a model of inherited NO deficiency. We showed dysfunction of glutathione metabolism in both ASL-deficient patients and *Asl*^*Neo/Neo*^ mouse model, whilst mRNA-LNP therapy corrected both ureagenesis and glutathione metabolism *in vivo*. We demonstrated the potential of [^18^F]FSPG-PET as a non-invasive diagnostic tool to assess the liver disease and therapeutic efficacy in ASA.

## Materials and Methods

### mRNA formulation

*hASL* and Luciferase (*Luc*) encoding mRNA encapsulated in Lipid Nanoparticles (LNPs) were provided by Moderna Therapeutics using their proprietary technology. Codon optimized mRNA encoding *hASL* was synthesized in vitro by T7 RNA polymerase-mediated transcription. The mRNA initiated with a cap, followed by a 5’ untranslated region (UTR), an open reading frame (ORF) encoding *hASL*, a 3’ UTR and a polyadenylated tail. Uridine was globally replaced with N1-methylpseudouridine, previously described (67). For in vivo intravenous delivery, LNP formulations were generated. Briefly, mRNA was mixed with lipids at a molar ratio of 3:1 (mRNA:lipid), previously described (31). mRNA-loaded nanoparticles were exchanged into final storage buffer and had particle sizes of 80 – 100 nm, >80% encapsulation of the mRNA by RiboGreen assay, and <10 EU/mL endotoxin levels.

### Reagents and antibodies

Full list of antibodies provided in **Table 1**, reagents in **Table 2** and instruments in **Table 3**.

**Table 1:** List of primary and secondary antibodies.

| Antibodies | Host | Company | Catalog num | Dilution used |
| --- | --- | --- | --- | --- |
| Anti-ACSL4 [F-4] | Rabbit | Santa Cruz Biotechnology | Sc-365230 | 1:200 WB |
| Anti-Actin | Rabbit | Cell Signalling | 4967 | 1:1000 WB |
| Anti-ASL | Rabbit | Abcam | Ab97370 | 1:1000 WB and histology |
| Anti-GAPDH | Mouse | Abcam | Ab97370 | 1:10000 WB |
| Anti-GPX4 | Rabbit | Abcam | Ab125066 | 1:500 WB |
| Anti-Nrf2 (fibroblast) | Rabbit | ThermoFisher | PA5-27882 | 1:500 WB |
| Anti-Nrf2 (liver) | Rabbit | Abcam | Ab62352 | 1:500 WB |
| Anti-GAPDH | Mouse | Abcam | Ab97370 | 1:10000 WB |
| Anti-xCT | Rabbit | Novus Biologicals | NB300-318 | 1:500 WB |
| Anti-Nitrotyrosine | Mouse | Merck Millipore | 05-233 clone 1A6 | 1:100 WB |
| Anti-GAPDH | Rabbit | Abcam | Ab9485 | 1:1000 WB |
| IRDye® 800CW goat anti-rabbit IgG |  | Licor | 926-32210 | 1:1000 in-cell<br>1:10,000 WB |
| IRDye® 680RD Donkey anti-mouse IgG |  | Licor | 923-68072 | 1:10,000 WB |
| CellTag™ 700 |  | Licor | 926-41091 | 1:1000 |
| HRP-linked anti-rabbit IgG |  | Cell Signalling | 7074 | 1:200 WB |

**Table 2:**
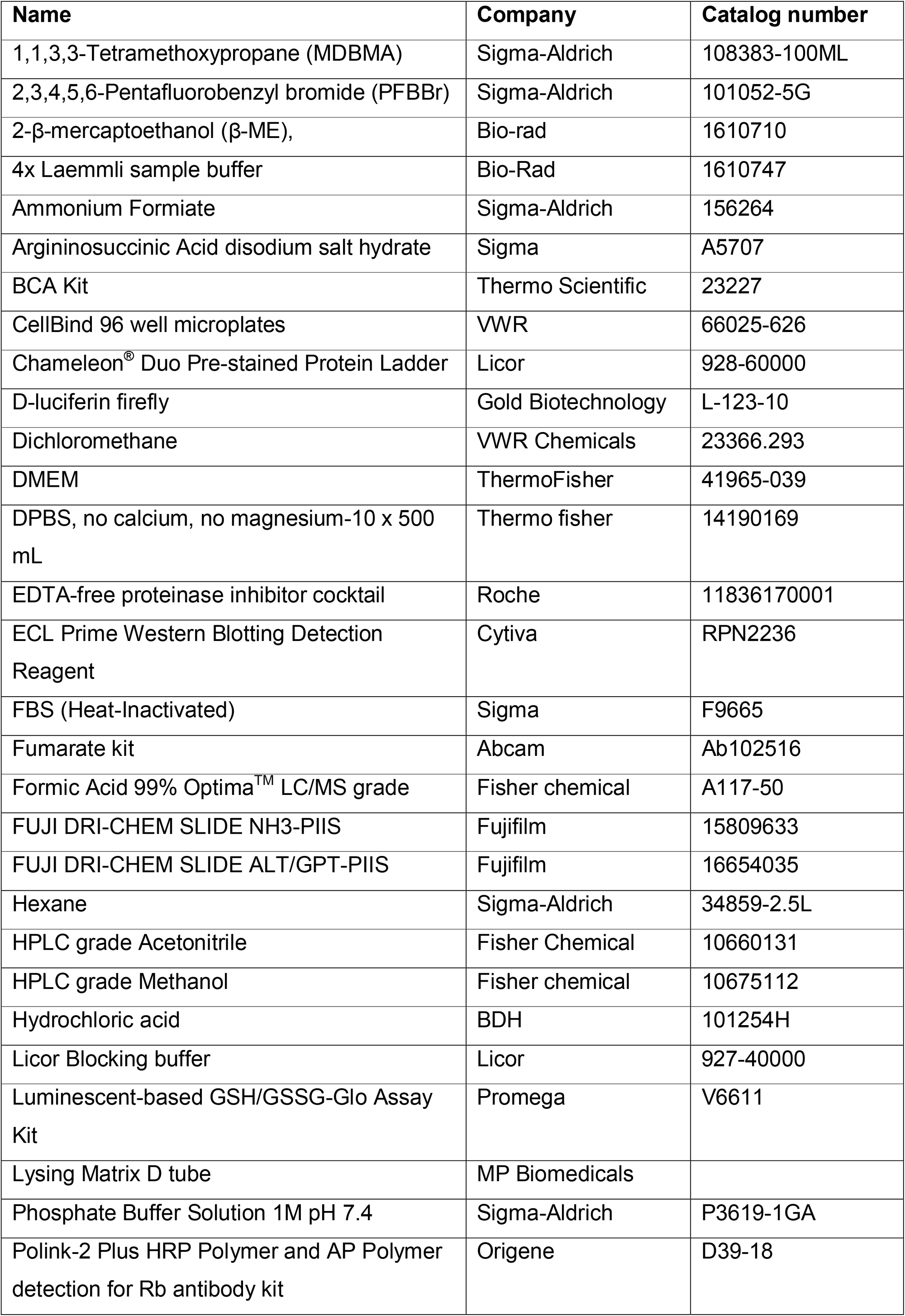
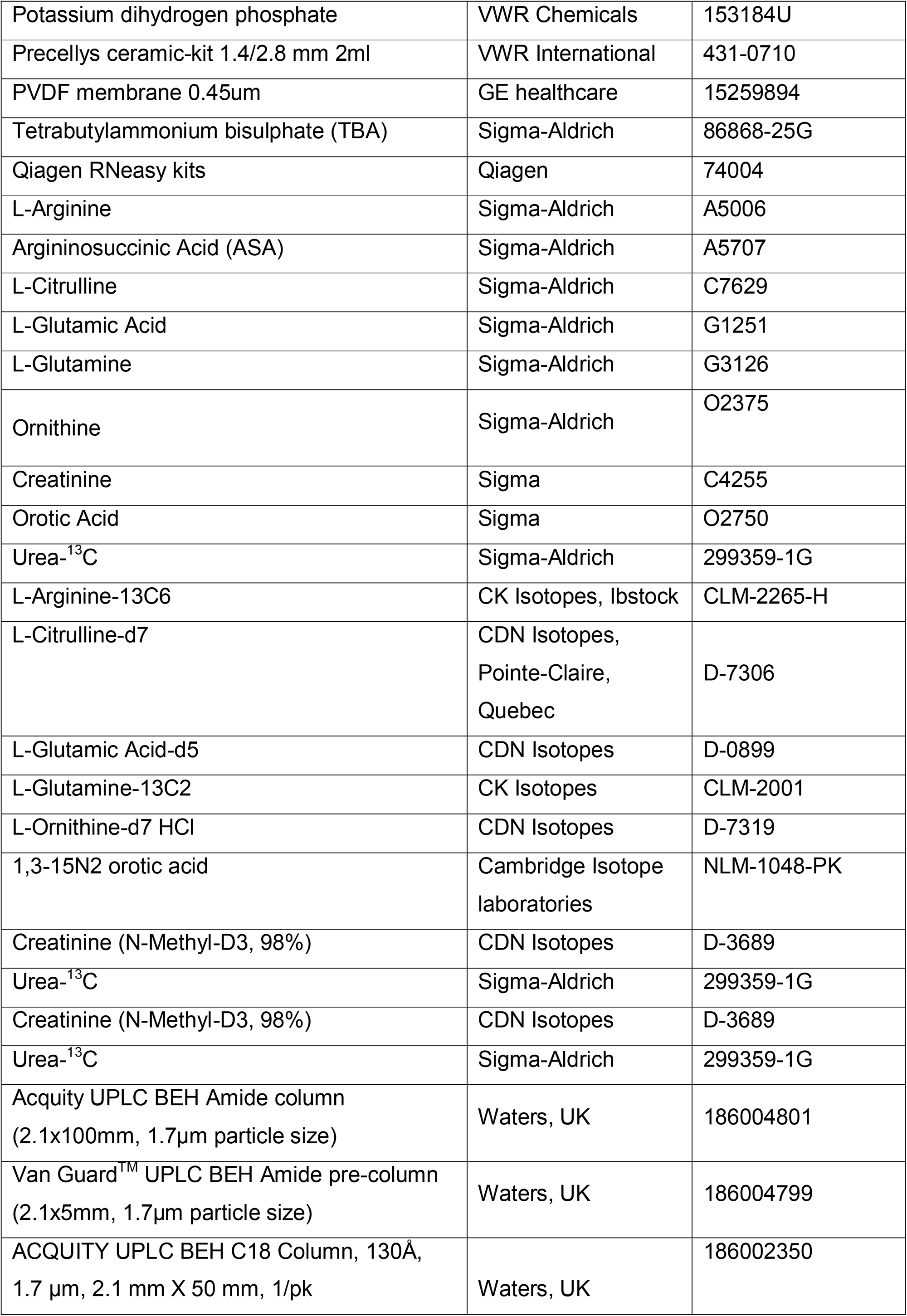
List of materials and reagents.

**Table 3:** List of instruments.

| **Instrument name** | **Company** |
| --- | --- |
| Capintech dose calibrator | Mirion medical |
| Concentrator plus | Eppendorf |
| DRI-CHEM NX600 | Fujifilm |
| FLUOstar Optima | BMG Labtech |
| GCMS Instrument | Thermoscientific |
| iBright™ Imaging System | ThermoFisher Scientific |
| IVIS® Spectrum In Vivo Imaging System | PerkinElmer, Waltham, US |
| In cell western Fluorescence Imaging scanner | Licor Odyssey CLx |
| NanoPET/CT plus system | Mediso |
| Precellys 24 tissue homogeniser | Bertin instruments |
| Precellys Evolution | Bertin Technologies |
| Xevo TQ-S | Waters, UK |
| Zeiss Axioplan | Zeiss |

### Patient Sample Study

Patients with ornithine transcarbamylase deficiency (OTCD), argininosuccinate synthase deficiency (ASSD) and argininosuccinate lyase deficiency (ASLD) followed at Great Ormond Street Hospital for Children NHS Foundation Trust, London, UK were asked to join a study with research ethics consent 13/LO/0168 issue by the Health Research Authority. Biological data were analysed anonymously.

### Animals

Animal procedures were approved by institutional ethical review and performed per UK home office licenses PP9223137, 70/14300 and PEFC6ABF1. *Asl*^*Neo/Neo*^ mice (B6.129S7-*Asl*^*tm1Brle*^/J) were purchased from Jackson Laboratory (Bar Harbor, ME) and maintained on standard rodent chow (Harlan 2018, Teklab Diets, Madison, WI) with free access to water in a 12-hour light / 12 hours dark environment. WT littermates were used as controls and housed in the same cages. Genotyping was performed using DNA extracted from tail clips as previously described (16).

### Animal experimental design

*Asl*^*Neo/Neo*^ animals were given systemic administration of *hASL* mRNA or *Luc* mRNA at dose of 1mg/kg or 2mg/kg for IV and IP injections, respectively. Untreated WT littermates were used as controls. For pharmacokinetics experiment, *Asl*^*Neo/Neo*^ mice were administered IV via tail-vein at 3-weeks of age and harvested at either 2, 24, 72 h or 7 days. In survival study for animals treated from birth, neonate pups at day 1 were administered mRNA intravenously through the temporal superficial vein, followed by intraperitoneal IP administration at day 7 at dose of 2 mg/kg and IV administration via tail-vein weekly from day 14 onwards up to 7-weeks of age. In survival study of animals treated from early adulthood, mice were given weekly IV administration through tail-vein from day 21 onwards up to 9-weeks of age. All harvests were performed 48 h following the last injection. Mutant mice were assigned randomly to study groups. All animals were monitored and weighted daily. In-life blood collection for longitudinal analysis was performed via tail-bleed and terminal blood collection via cardiac puncture, followed by cervical dislocation for harvest. Urine samples were collected longitudinally and post-harvest on Whatman filter paper. Animals were culled either at the stated endpoint or at the occurrence of >15% body weight loss and/or reaching severity limit.

### [^18^F]FSPG PET imaging

[^18^F]FSPG was synthesized using a GE FASTlab automated synthesis module and quality control performed as previously reported (68). Anaesthetized (1.5-2% isofluorane in O_2_) 2-3 week-old *Asl*^*Neo/Neo*^ mice with age-matched WT littermates were imaged following IV injection of 1-3 MBq of radiotracer. For rescue experiment with mRNA therapy, *Asl*^*Neo/Neo*^ mice were given 1 mg/kg IV dose of *hASL* mRNA dose at day 1 of birth followed with weekly 2 mg/kg IP injection. Age-matched control untreated *Asl*^*Neo/Neo*^ and WT littermates were included. For all animals, dynamic PET scans were acquired between 40 and 90 min post-injection on a Mediso NanoScan PET/CT system (1-5 coincidence mode; 3D reconstruction; CT attenuation-corrected; scatter corrected) using a four-bed mouse hotel (Mediso) (69). CT images were acquired for anatomical visualization (360 projections; helicalacquisition; 55 kVp; 600 ms exposure time). A dynamic iterative reconstruction algorithm, Tera-Tomo 3D (0.4 × 0.4 × 0.4 mm^3^ voxel size), was used with attenuation, scatter, and random coincidences correction. Radiotracer concentration was quantified using VivoQuant software (v 2.5, Invicro Ltd), with volumes of interest drawn manually using the CT image as reference. Data were expressed as percent injected dose per gram of tissue (% ID/g). Following [^18^F]FSPG PET, mice were culled by cervical dislocation and liver, skin and other tissue was collected, snap frozen and moved to a -80°C freezer for later *ex vivo* analysis.

### Ammonia and Alanine aminotransferase (ALT) measurement

To obtain plasma samples, whole blood was collected in EDTA tube (Sarstedt, Germany) and centrifuged immediately at 13,000 rpm for 5 min at room temperature. Supernatant was then transferred into a microcentrifuge tube and stored at -80^°^C. Ammonia and ALT reads were obtained from Fujifilm NX600 machine using ammonia and ALT cartridges respectively (Fujifilm, Japan) using 10μl plasma volume (diluted vol:vol 1:3 in PBS).

### Amino acid analysis

Liquid chromatography-Mass spectrometry (LC-MS/MS) was used for amino acid measurements (argininosuccinic acid and L-citrulline) from dried bloodspots as described previously (11) using the hydrophilic interaction liquid chromatography (HILIC) separation of metabolites, method adapted from (70). Briefly, 40μl of whole blood was spotted on Guthrie blood spot card, dried at room temperature for 24h and stored in -20^°^C in a foil bag with desiccant. 3mm blood spot punch was extracted in 100μl methanol containing stable isotopes (2nmol/l, L-citrulline-d7, CDN isotopes, Pomite-Claire, Quebec), used as internal standards, for 15 min in sonicating waterbath at room temperature. The supernatant was collected and dried using Eppendorf® Concentrator Plus and resuspended in 80μl of 0.05M HCl, topped with 280μl of Solvent A (10mM ammonium formiate+85% Acetonitrile (ACN)+0.15%Formic acid (FA)), centrifuged at 16,000rpm for 5 min and supernatant taken for analysis.

Acquity UltraPure Liquid Chromatography (UPLC)-system (Waters, Manchester, UK) using Acquity UPLC BEH Amide column (2.1×100mm, 1.7μm particle size) and Van Guard^™^ UPLC BEH Amide pre-column (2.1×5mm, 1.7μm particle size) (Waters Limited, UK) was used for amino acid chromatography. The mobile phases were (A) 10mM ammonium formiate in 85% ACN and 0.15% FA and (B) 15mM ammonium formiate containing 0.15% formic acid, pH 3.0. Detection was performed using a tandem mass spectrometer Xevo TQ-S (Waters, Manchester, UK) using multiple reaction monitoring in positive ion mode. The dwell time was set automatically with MRM-transition of 291.2>70.2, 273.2>70.2 and 176.1>159 respectively for ASA, ASA-anhydrides and L-citrulline. L-Citrulline-d7 (183.15>166.05) was used as internal standard control. Argininosuccinate data were analysed using Masslynx 4.2 software (Micromass UK Ltd, Cheshire, UK) and TargetLynx^™^ application manager used for subsequent batch analysis.

### Nitric oxide metabolites

Plasma samples were pre-treated with N-ethylmaleimide (NEM) and deproteinised by precipitation with methanol (v:v 1:1), followed by centrifugation at 16,000c×g for 20cmin. Liver samples were diluted 1:3 (w/v) with homogenisation solution (10mM PBS supplemented with 10mM NEM and 2.5mM EDTA) and homogenised on ice using an all-glass Kimble tissue grinder in combination with a GlasCol GT Series stirrer (8 up-and down strokes). Tissue homogenates were deproteinized in the same manner as plasma. Deproteinized samples were analysed for nitrate (NO_3_ ^-^) and nitrite (NO_2_ ^-^) using a dedicated high-performance liquid chromatography analyser (ENO20, Eicom) as described (71).

Tissue homogenates were analysed for content of total nitrosation products (RXNO) by gas-phase chemiluminescence of bound NO following reductive denitrosation. Nitrite was removed from sample aliquots by addition of 10% (v:v) of a reaction solution comprising 5% sulfanilamide in 1 M HCl and reaction for 15 min prior to injection into an acidic triiodide-containing reduction chamber. The amount of NO liberated from low-molecular weight and protein nitroso-species was quantified by a gas-phase chemiluminesence analyser (CLD 77cam sp, EcoPhysics), as previously described (71).

### Oxidative stress marker and thiol redox status

Circulating lipid peroxidation products in plasma were evaluated by measuring the thiobarbituric acid reactive substances (TBARS) essentially as described elsehwere (72). Malondialdehyde (MDA), a major breakdown product of the peroxidation of unsaturated fatty acids, in plasma or liver homogenate reacts with thiobarbituric acid, at high temperature and acidic conditions, to form a coloured adduct with maximum absorption at 532 nm. After subtraction of background coloration, the resultant absorbance at 532 nm in the sample is then compared to that of a standard curve of solutions of known concentrations of MDA.

Thiol redox status in plasma and liver homogenates was measured using ultra-high performance liquid chromatography tandem mass spectrometry (UPLC-MS/MS) following derivatization with the thiol alkylans NEM, as described in detail elsewhere (73). The LC-MS system was used to separate and quantify the biological aminothiols including total glutathione (GSH and GSSG), cysteine (CyS), cystine (CySS), homocysteine (HCyS), homocystine (HCySS), glutamyl-cysteine, cysteinylglycine as well as sulfide. In addition to the free thiols in the sample, their total concentrations (free + protein-bound forms and disulfides) were determined after sample pre-processing with dithiothreitol (DTT). For this purpose, aliquots of plasma and tissue homogenates already reacted with NEM were subjected to reduction by addition of 50mM DTT (1:1 v:v). Following incubation for 30cmin at room temperature for complete reduction, excess NEM (100mM; 1:10, v:v) was added for derivatization of the liberated thiols and samples were processed as before. NEM-derivatized sample aliquots were spiked with stable-isotope labelled internal standards, subjected to ultrafiltration for protein removal and diluted in 10 mM ammonium phosphate buffer before analysis by LC-MS/MS.

### ASL enzyme activity

For liver ASL activity, 20-30mg of liver was homogenised in 400μl of cold homogenising buffer (50mM phosphate buffer pH 7.5 and 1x Roche EDTA-free protease inhibitor (Roche, Switzerland)) using Precellys homogeniser tube (VWR, UK) and Precellys 24 tissue homogeniser (Bertin Instruments, France), centrifuged at 10000g for 20 min at 4^°^C and protein levels measured from the supernatant using BCA kit (Thermo Fisher Scientific, UK). 60μg of protein lysate was incubated with 3.6mM ASA in final volume of 50μl, incubated at 37^°^C for 1h followed by reaction termination at 80^°^C for 20 min. The mixture was centrifuged at 10000g for 5 min and 5μl of the supernatant was used to measure fumarate levels per instruction from the commercial fumarate kit (Abcam, Cambridge, UK).

For ASL enzymatic assay in fibroblasts, 800,000 cells were plated 24h before in a 6cm tissue culture grade dish. 2.5μg of hASL mRNA-LNPs or Luc mRNA-LNPs were transfected and incubated for 48h. Cells were harvested in 250μl of assay buffer provided in fumarate kit. 60μg of protein per samples was incubated with 300μM of ASA in final volume of 100μl, incubated for 15 min at 37^°^C followed by reaction termination at 80^°^C for 15 min. Samples were then centrifuged at room temperature for 5 min at maximum speed on benchtop centrifuge. 50μl of supernatant was used for fumarate reaction to determine fumarate reaction per the kit instructions.

### Orotate measurement

Urine was spotted on a Whatman filter paper and dried over 24h room temperature and stored in -20^°^C in a foil bag with desiccant, method adapted from (74). For extraction, a 3mm punch of the urine was eluted in 150μl of ddH_2_O containing 40μM labelled orotic acid (stable isotope 1,3-15N2 orotic acid, Cambridge isotopes, Andover, MA) and creatinine-D3 (N-Methyl-D3, CDN isotopes) at room temperature for 3h. Orotic acid was analysed in negative ion mode using LC-MS/MS on waters Xevo-TQ-S with MRM transitions (155.1>111.1 and 157.1>113.1) respectively for labelled and unlabelled orotic acid. An isocratic method was used with sample eluted at flow rate of 0.25ml/min with 40% solvent A (water +0.1% FA) and 60% ACN for 1.5 min followed by wash with 100% ACN for 0.5 min and 0.5 min of initial starting condition. Creatinine was used as control to normalise orotic acid levels from the same extracted sample. Creatinine was measured in positive ion mode with MRM transitions (113.95>43.85 and 116.95>46.85) for unlabelled and labelled creatinine respectively with same LC-MS/MS conditions as orotic acid.

### Cell culture

Fibroblasts cells were maintained in Dulbecco’s modified Eagle medium (ThermoFisher Scientific, 41965062) supplemented with 10% (vol/vol) Fetal Bovine Serum (Sigma-Aldrich, F9665) and 50 units of Penicillin and Streptomycin (ThermoScientific, P4458) and maintained at 37^°^C in a humidified 5% CO_2_-air atmosphere. Healthy control fibroblast line was obtained commercially (Lonza, CC-2511). ASL deficient patient fibroblasts were obtained from 2 patients who joint a study with research ethics consent 13/LO/0171 issue by the Health Research Authority. The genotype of patient 1 was c.437G>A / c.437G>A; R146Q / R146Q. The genotype of patient 2 was c.719-2A>G / c.857A>G; ? / GlN286Arg.

### In-cell western

Fibroblasts were plated at a density of 5,000 cells per well in a 96 well plate (CellBind 96 well microplates, 66025-626). The next day cells were transfected with 0.2μg of *hASL* mRNA or *Luc* mRNA per well for 24 hours after which the cells were fixed in ice-cold methanol for 15 minutes. Rest of the procedure was performed in room temperature. Following 3 quick washes with 1xPBS, the wells were blocked with Licor blocking buffer (927-40000, Licor, Cambridge, UK) for 90 minutes followed by incubation with Anti-ASL antibody (Abcam) for 2 hours. The wells were washed 3x with 1xPBS, incubated with anti-rabbit secondary (IRDye® 800CW Goat anti-Rabbit IgG 1:1000, 926-32210, Licor) and cell dye (CellTag700, 1:1000, 926-41091, Licor) for 1h and washed 3x with 1xPBS for 5 minutes after. Post final wash, PBS was removed, plates dried and read on Licor Odyssey CLx (settings: 4mm focus offset, lowest setting quality, resolution 169μM, 700 and 800 channel). Acquisition and analysis were performed in the Licor ImageStudio Lite software (Licor, Cambridge, UK).

### *Ex vivo* total glutathione quantification

Frozen liver and skin tissue was thawed on ice and 25-50 mg of tissue was added to prechilled Lysing Matrix D tube (MP Biomedicals) containing ice cold 400 μL 1X passive lysis buffer (Promega; E1941). The tissue was then lysed at 4 °C on a Precellys Evolution (Bertin Technologies, Montigny-le-Bretonneux, France); samples run for five 30s cycles at 6700 RPM. Lysates were centrifuged at 15,000 × *g* for 10 min at 4°C and the supernatant taken for analysis. Total intracellular glutathione was determined using the luminescent-based GSH/GSSG-Glo Assay Kit (Promega; V6611) according to manufacturer’s instructions in a white 96-well plate prepared with 5 μL of sample supernatant (neat or 1:10 diluted) along with 5 μL of GSH standards (1-100 μM). Results were normalized to protein concentration, determined using the Pierce™ BCA Protein Assay Kit (ThermoFisher Scientific, city, country) as per the manufacturer’s instructions.

### Western blot

Western blot analysis for was carried out using the iBind Flex system (ThermoFisher Scientific), a previously published method (26), for antibody immunoblotting. To produce lysates frozen liver and skin tissue was processed and the protein concentration determined as above, except, the prechilled ice cold 400 μL RIPA buffer, with 1% Proteinase and phosphatase inhibitors (HALT) was used instead of 1X passive lysis buffer.

Membranes were probed using rabbit polyclonal anti-xCT (1:500; Novus Biologicals; NB300-318). Actin was used as a loading control for all experiments (1:1000; Cell Signaling Technology; 4967). HRP-linked anti-rabbit IgG (1:200, Cell Signalling Technology; 7074) was used as secondary antibody.

Protein bands were visualized using ECL Prime Western Blotting Detection Reagent (Cytiva; RPN2236) as per the manufacturer’s instructions and the iBright™ Imaging System (ThermoFisher Scientific). Image analysis and band quantification was performed using the iBright™ Analysis Software (ThermoFisher Scientific). xCT protein signal per sample was normalised against actin which was used as a loading control.

For ASL and nitrotyrosine levels analysis in liver, 30mg of liver was homogenised in ice-cold 1x RIPA buffer (Cell Signalling) using Precellys homogenising tube and homogeniser, centrifuged at 10000g for 20 min at 4^°^C and protein levels measured from the supernatant using BCA kit. 40μg of protein per sample was diluted 1:1 with 2x Laemmli sample buffer (containing 10% 2-β-mercaptoethanol (β-ME)) at final volume of 40μl, vortexed and heated to 95^°^C for 10 min. SDS-PAGE was used to separate the proteins at 100V for 1h followed by wet transfer of proteins into an immobilin PVDF membrane at 400mA for 1h. The membrane was blocked in 5% non-fat milk powder in PBS-T (1xPBS with 0.1% tween-20) followed by overnight incubation at 4^°^C with primary antibodies (Anti-ASL, Abcam ab97370, 1:1000; Anti-GAPDH mouse, Abcam ab8245, 1:10,000; Anti-nitrotyrosine, Merck 05-233, 1:100; Anti-GAPDH rabbit, Abcam ab9485, 1:1000), 3x 5-min washes with PBS-T, 1h incubation with fluorescent secondary antibodies (IRDye® 800CW Goat anti-Rabbit IgG 1:1000, 926-32210 and IRDye® 680RD Donkey anti-Mouse IgG, 923-68072, Licor) and 3x 5 min washes with PBS-T. Image acquisition and analysis was performed using Licor Odyssey and image analysed using Licor ImageStudio Lite software. ASL protein signal per sample was normalised against GAPDH which was used as a loading control.

### Histology

At harvest, liver was fixed in 10% formalin solution, left at room temperature for 48h before transferring and storing in 70% ethanol. The liver was paraffin embedded and sectioned at 5μM thickness. Sections were dewaxed in histoclear, hydrated through graded ethanol solution to water followed by incubated in 1% H_2_O_2_ to remove blood stains. Antigen retrieval was performed in boiling 0.01M citrate buffer for 20 min and then cooled to rt. The slides were blocked in 15% goat-serum and TBST-T for 30 min in rt then incubated overnight with primary antibody 9Anti-ASL, ab97370, Abcam, 1:1000) in 10% goat serum and washed 3x with TBST-T. DAB staining was performed using Polink-2 Plus HRP Polymer and AP Polymer detection for Rb antibody kit (D39-18, Origene) following the manufacturer’s instructions. The slides were then dehydrated with increasing gradient of ethanol to water and histoclear. The slides were mounted using non-aqueous mounting media – Microscopy DPX (Merck) and dried overnight.

The slides were imaged under Zeiss Axioplan Histology scope at UCL Great Ormond Street Institute of Child Health Imaging Facility. 10 images per condition was taken in random and averaged. Images were analysed using Fiji software using macro written by Dr. Dale Moulding from UCL Great Ormond Street Institute of Child Health Imaging Facility. The macro utilises colour deconvolution to quantify DAB percentage coverage to calculate percentage of ASL positive regions.

### ^13^C ureagenesis

30 min pre-harvest animals were given intraperitoneal (IP) administration of 1% body weight labelled sodium acetate (1,2-^13^C_2_, 99%, CLM-440-1, CK Isotopes). Plasma was harvested as before and stored at -80^°^C until analysis. Samples were processed and analysed usingisotope-ratio mass spectrometry. Mouse plasma (25 μL) was deproteinized by addition of 25μL of 60% perchloric acid and 0.5 mL 5mM urea (added as unlabelled carrier), sample vortexed, and centrifuged at 21130*g* for 5 minutes to remove precipitated protein. The tube was then left uncapped for 30 minutes at room temperature to facilitate evaporation of CO_2_. The supernatant was transferred to a new microcentrifuge tube and 100μl 0.5M potassium phosphate added. The pH was adjusted to 4 –7 with 1M KOH solution using pH strips. Sample centrifuged at 21130*g* for 5 minutes to remove the precipitated potassium perchlorate. The supernatant was added to an ion exchange column (1ml Dowex-1 1×8-200 resin in empty polypropylene SPE Tube), and eluant collected into a 12ml Exetainer (Labco. UK Ltd). The ion exchange column was washed with 2mL 10 mM HCl, and the eluant combined with the earlier fraction in the Exetainer. Samples were dried under N_2_ at 80°C. Dried samples were left uncapped in sealed dessicator for 18 hours with gauze soaked in 1M NaOH to absorb any residual trace of bicarbonate/CO_2.._ The tube was sealed with a cap, and flushfilled with 75ml/min helium for 5 minutes per tube. Meanwhile, 8ml 0.5 M potassium phosphate, pH 6.0 was heated to boil for 5 minutes to remove dissolved gas, then cooled. After cooling, 60mg urease dissolved in buffer by very gentle vortexing. 400 μL urease in potassium phosphate buffer injected through the septum of each vial using a gastight syringe, avoiding introduction of any air bubbles. Samples incubated for 60 minutes at 25°C and 100μL 20% phosphoric acid through the septum, using a gastight syringe, avoiding introduction of any air bubbles. Samples incubated for a further 60 minutes at 25°C to allow full release of CO_2_. Samples analysed by Thermo-Finnigan DeltaPlus XP Plus isotope ratio-mass spectrometer (Thermo Fisher Scientific, Bremen, Germany) with Gasbench sample Introduction unit and CTC GC-PAL autosampler, with 10 technical replicates (final 10 of 15 sample injections of 100μL). Urea production was calculated from the ^13^CO_2_/^12^CO_2_ ratio, taking into account the initial dilution by carrier urea, and the concentration of urea in the plasma samples.

### Bioluminescence imaging

Animals were anesthetized with isoflurane (Abbott Laboratories, Illinois, US), injected intraperitoneally with D-luciferin firefly (15mg/ml in PBS) (L-123-10, Gold Biotechnology, Olivette, US) at a dose of 150mg/kg and imaged 5 min later with a cooled charge-coupled device (CCD) camera in the IVIS^®^ Spectrum in vivo imaging system (IVIS; PerkinElmer, Waltham, US). Grey-scale photographs were acquired with a 24-cm field of view and then a bioluminescence image was obtained using a binning resolution factor of 4, a 1.2/f stop and open filter. Regions of interest (ROIs) were defined manually using a standard area for the mouse liver. Signal intensities were calculated with Living Image software (Perkin Elmer, Waltham, US) and expressed as photons per second per cm^2^ per steradian. At each timepoint, bioluminescence imaging was carried out with PBS injected control rodents to establish a median baseline; data points were expressed as fold-change over this internal standard for each individual animal.

### Transcriptomics

RNA was extracted from liver samples using Qiagen RNeasy kits (74004) following kit instructions. Liver samples from the WT, Luc mRNA and hASL mRNA neonatal treated group of *Asl*^*Neo/Neo*^ mutants were analysed. In all cases, cDNA libraries were prepared using the Kapa mRNA Hyper Prep kit (KapaBiosystem, Pleasanton, CA, USA) according to the manufacturer’s instructions, and sequenced on an Illumina NextSeq 1000/2000 to generate ∼16 million 50-bp paired end reads per sample (UCL Genomics). Fastp was used for adapter trimming, read filtering and base correction. Processed reads were mapped to the GRCm38 mouse reference genome via STAR using gene annotations from GENODE M29. Normalization and differential gene expression analyses were carried out using the DESeq2 R package (v2.12) (75) with differentially expressed genes (DEGs) defined on the basis of a log2-fold change > 0.1 or < -0.1, and an FDR-corrected p-value of < 0.05. Volcano plots were generated to visualize the results of analyses between *Luc* mRNA against WT, *hASL* mRNA against WT, and *hASL* mRNA against *Luc* mRNA. The differential expression of specific genes of interest was plotted in a heatmap with their respective molecular pathways. The data were visualized using ggplot2R R package (v3.3.5) (76).

### RT-PCR of GCL and GS

Liver samples were stored frozen at −80 °C before RNA extraction with the RNeasy kit (QIAgen, Crawley, UK) according to the manufacturer’s instructions. cDNA was amplified using High-Capacity RNA-to-cDNA™ Kit (Applied Biosystems, Rockford, IL, USA). The GCLC sequence was amplified using the following primers: 5′-ACACCTGGATGATGCCAACGAG-3′ (forward) and 5′-CCTCCATTGGTCGGAACTCTAC-3′ (reverse), the GCLM sequence was amplified using the following primers: 5′-TCCTGCTGTGTGATGCCACCAG-3′ (forward) and 5′-GCTTCCTGGAAACTTGCCTCAG -3′ (reverse) and the GS sequence was amplified using the following primers: 5′-CCAGGAAGTTGCTGTGGTGTAC-3′ (forward) and 5′-GCTGTATGGCAATGTCTGGACAC-3′ (reverse). Amplification was detected and normalised against Titin which was amplified using the following primers: 5′-AAAACGAGCAGTGACGTGAGC-3′ (forward) and 5′-TTCAGTCATGCTGCTAGCGC-3′ (reverse). Amplification reactions were carried out using 5 μl of sample, 2.5 μmol.l−1 of each primer, and SYBR green master mix using the Luna universal qPCR master mix (New England Biolabs, Ipswich, USA) for a 25 μl reaction. The amplification conditions were 95 °C for 10 min followed by 40 cycles of 95°C for 15s, 60°C for 45s. Data were processed with StepOneTM software (ThermoFisher Scientific, Rockford, IL, USA).

### Statistical analysis

Data was analysed and represented using Graphpad Prism 9.0 software (San Diego, CA, USA). Figures shown mean +/-standard deviation. Kaplan-Meier survival curves were analysed using log-rank test. Student’s t-test was performed to compare two groups and one-way ANOVA was used to compare more than 2 groups with Tukey’s post-test comparison. The groups passed normality test. Log transformation was used to compare groups lacking normal distribution.

## Author Contributions

THW and JB designed the study. SG, OVT and DP conducted most of the experimental work. LT, YK, ARB, RSE, PBM, SNW and PG contributed to technical assistance in experimental work. ALGM and MR analysed the transcriptomics data. MM and MF analysed the thiol reactome data. SF and MM collected patients’ data. SE analysed the ureagenesis with stable isotope. AC, SS, LR, PGVM, AF provided the *Luc* and *hASL* mRNA constructs. SG wrote the manuscript. All authors reviewed and approved the manuscript.

## Acknowledgements

The authors thank Phil Blower, Kavitha Sunassee, Jana Kim, Floyd Laniyan, and the staff from Biological services for help with PPL licence, breeding and maintenance of the ASL colony at King’s College London and University College London. The authors thank the patients and families for participating in the study.

## Supplementary figures

**Supplementary Figure 1.**
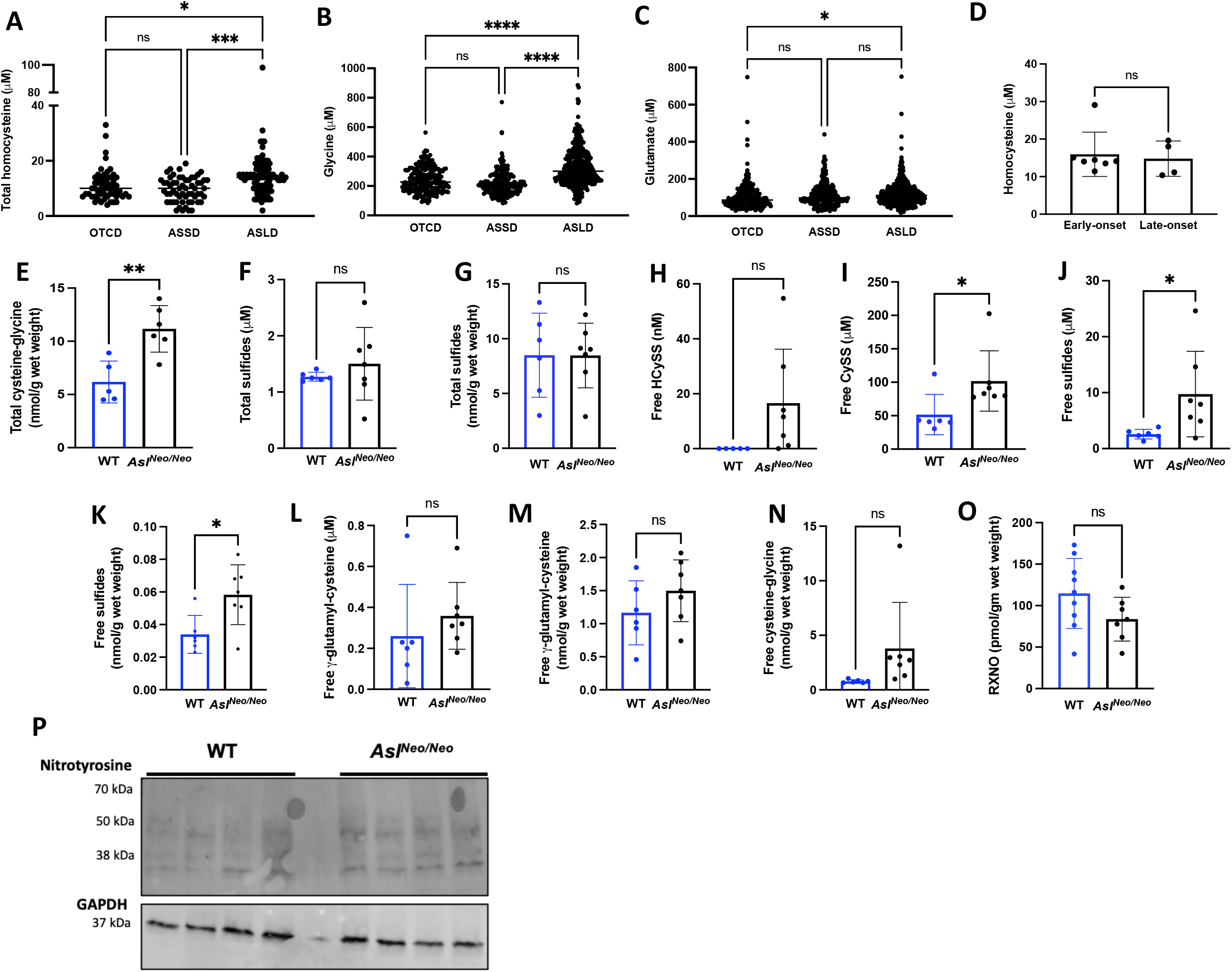
Dysfunction of glutathione metabolism in ASL-deficient patients and mouse model *Asl*^*Neo/Neo*^. (**A**) Collated single measurements of plasma total homocysteine, (**B**) glycine and (**C**) glutamate of patients followed for OTCD, ASSD and ASLD. (**D**) Mean plasma total homocysteine levels between early- and late-onset ASLD. (**E**) Liver cysteine-glycine, (**F**) plasma and (**G**) liver total sulfides in 2-weeks old *Asl*^*Neo/Neo*^ versus WT littermates. (**H**) Plasma homocysteine, (**I**) plasma cystine, (**J**) plasma and (**K**) liver sulfides, (**L**) plasma and (**M**) liver free γ-glutamyl-cysteine and (**N**) liver free cysteine-glycine in 2-weeks old *Asl*^*Neo/Neo*^versus WT littermates. (**O**) Decreased trend of nitroso-species (RXNO), including N-nitrosospecies (RNNO) and S-nitrosospecies (RSNO) in *Asl*^*Neo/Neo*^ versus WT littermates. (**P**) Liver nitrotyrosine levels by western blot between *Asl*^*Neo/Neo*^ mice and WT. (**A-C**) One-way ANOVA with Tukey’s post-test (**D-O**) Unpaired two-tailed Student’s t test; * p<0.05, ** p<0.01, *** p<0.001, ns not significant. **(A)**: OTCD n=60, ASSD n=59, ASLD n=97. **(B, C)**: OTCD n=220, ASSD n=198-199, ASLD n=352-353. **(D)**: Early-onset ASLD n=7, late-onset ASLD n=4. (**E-O**) WT n=6-8; *Asl*^*Neo/Neo*^ n=7. ASSD: argininosuccinate synthase deficiency; ASLD: argininosuccinate lyase deficiency; CySS: cystine; GSH: glutathione; HcyS: homocysteine; HcySS: homocystine; OTCD: ornithine transcarbamylase deficiency. Graphs show means +SD.

**Supplementary Figure 2.**
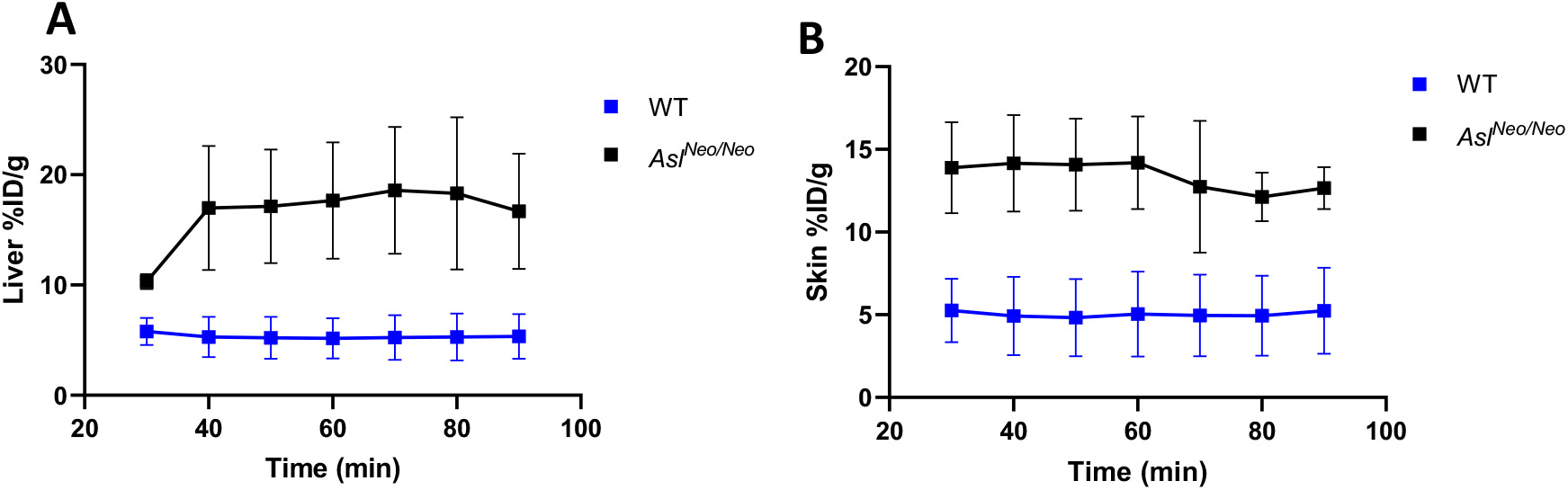
[18] FSPG radiotracer PET scan in *Asl*^*Neo/Neo*^ mice. [^18^F] FSPG PET in Asl^Neo/Neo^ and WT mice showed that **(A)** liver and **(B)** skin [^18^F] FSPG concentration (%ID/g) was seemingly unchanged between 40 and 90 minutes post injection.

**Supplementary Figure 3.**
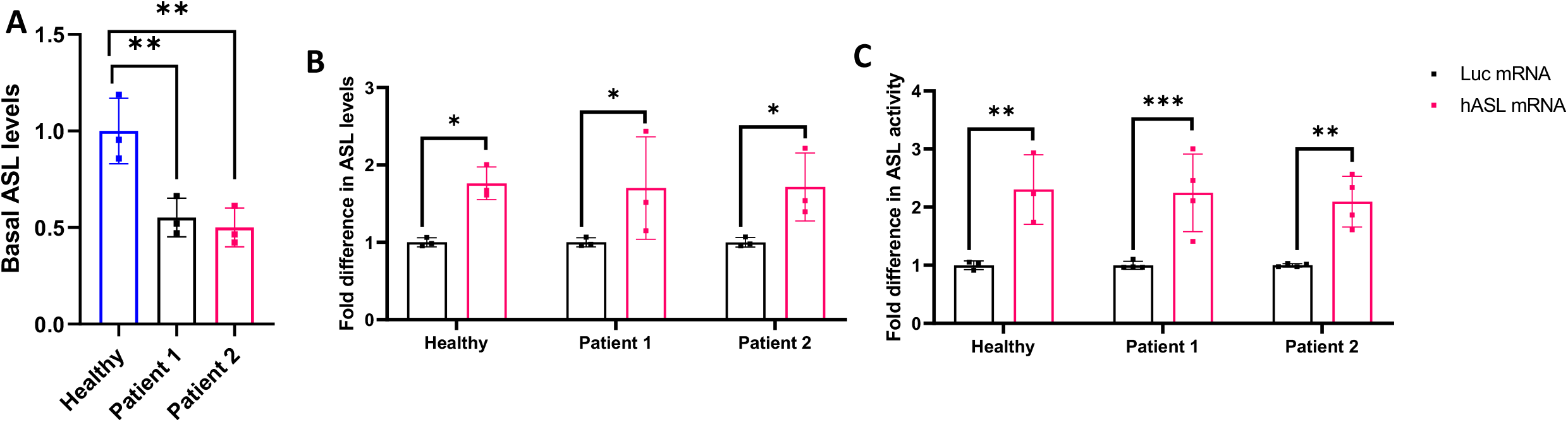
*In vitro* efficacy of *hASL* mRNA. (**A**) Basal expression levels of ASL in fibroblasts from control and two ASA patients. Fold difference in (**B**) ASL levels and (**B**) ASL activity following 24 hours and 48 hours incubation with either *Luc* mRNA or *hASL* mRNA, respectively from 3 independent experiments and normalised to healthy control (**A**) and Luc mRNA treated control (**B, C**). Statistical analysis by One-way ANOVA with Dunnett’s multiple comparisons test against healthy control (**A**) and two-way ANOVA with uncorrected Fisher’s Least Significant Difference (LSD) (**B, C**), * p<0.05, ** p<0.01, *** p<0.001. Graphs show means +SD.

**Supplementary Figure 4.**
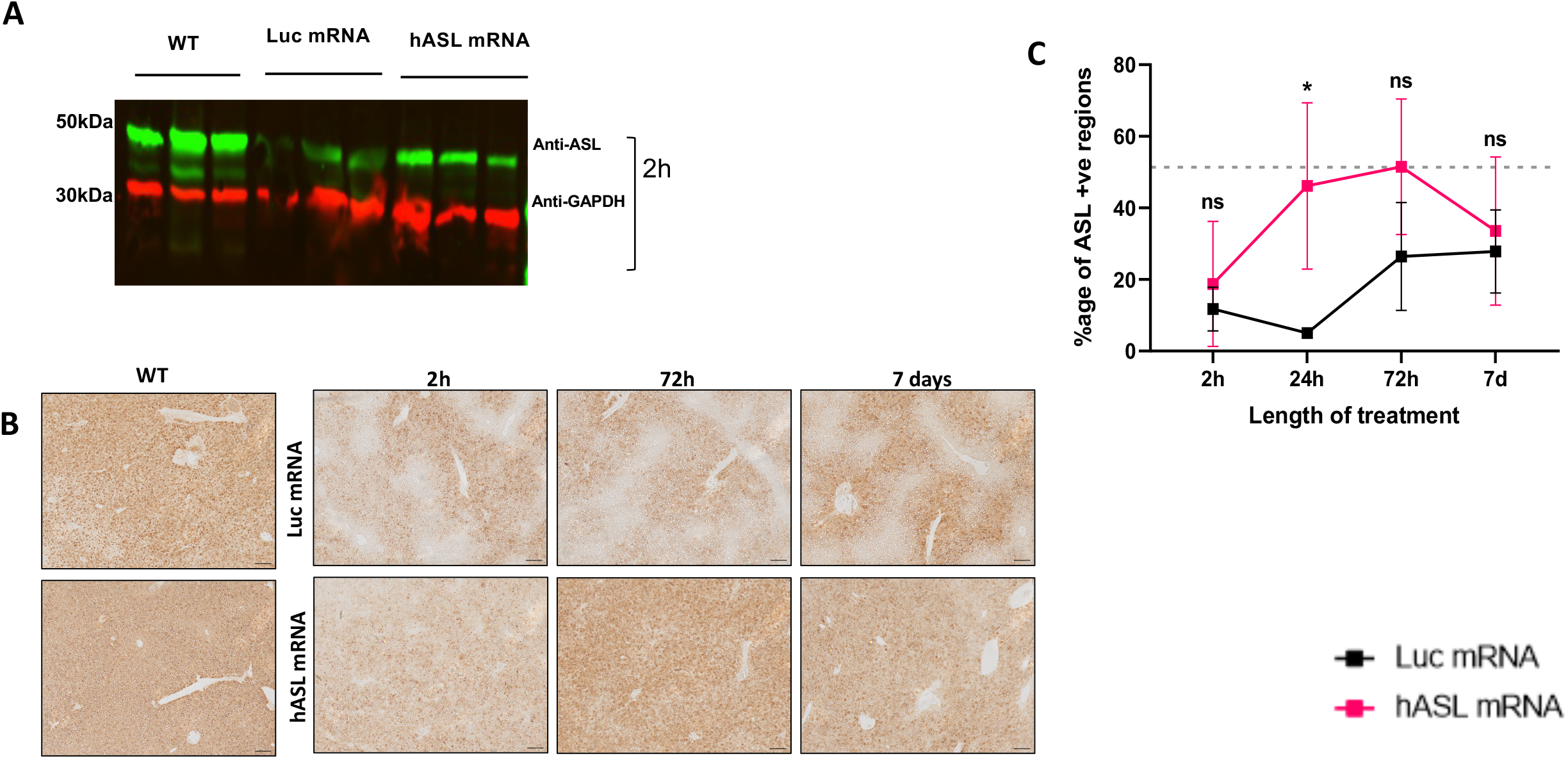
Pharmacokinetics of *hASL* mRNA in *Asl*^*Neo/Neo*^ mice. (A) ASL western blot at 2 hours post mRNA administration (n=3). (**B**) Representative images of liver ASL immunostaining at 2 hours, 72 hours and 7 days post mRNA administration from WT and *Luc* mRNA or *hASL* mRNA treated *Asl*^*Neo/Neo*^ mice and (**C**) quantification normalized to WT scaled to 1 (grey dotted line). (**B**) Scale bar= 100μM. One-way ANOVA with Tukey’s post-test per timepoint, ns=not significant, *p<0.05, n=3. Graph show means +SD.

**Supplementary Figure 5.**
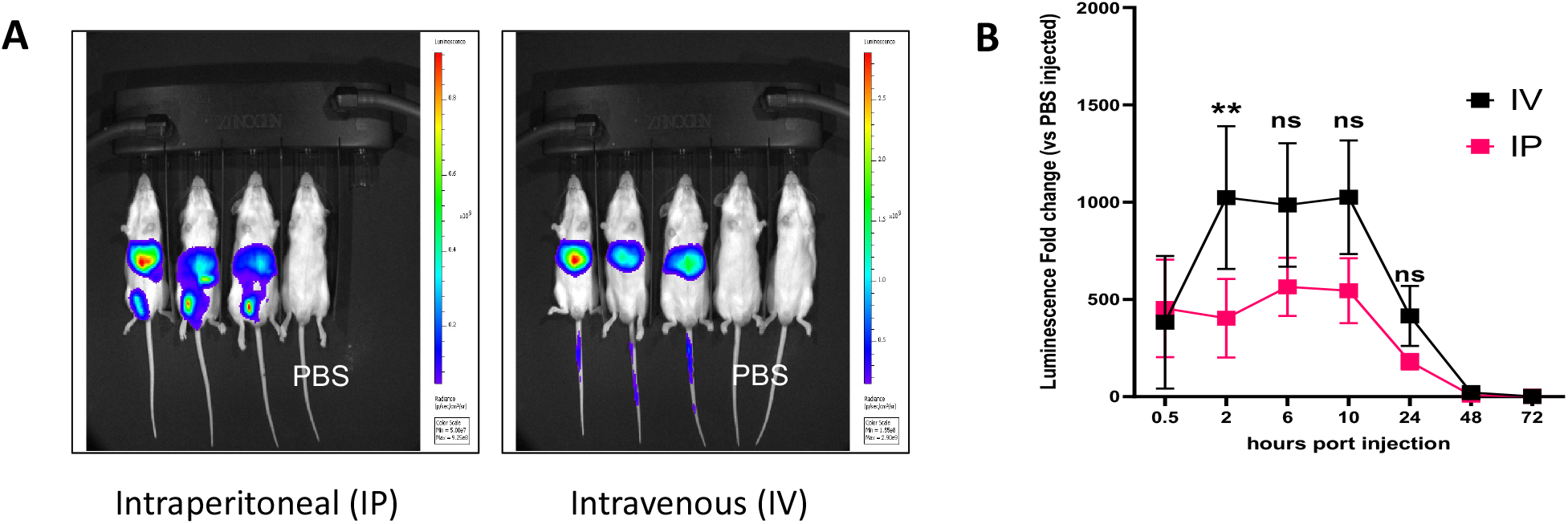
Equivalence of liver biodistribution between intravenous and intraperitoneal administration. **(A)** Representative *in vivo* luminescence image of WT CD1 strain animals injected with either PBS (IP) or *Luc* mRNA either intraperitoneally or intravenously 10 hours post injection. **(B)** Luminescence fold change in liver post 0.5, 2, 6, 10-, 24-, 48- and 72-hour injection, show two-fold difference in efficacy between IV vs IP. Readings normalised to PBS injected controls. Unpaired two-tailed t-test per timepoint, ns= not significant, **p<0.01, IV and IP n=3, PBS n=4. Graph show means +SD. IV = intravenous, IP= intraperitoneal.

**Supplementary Figure 6.**
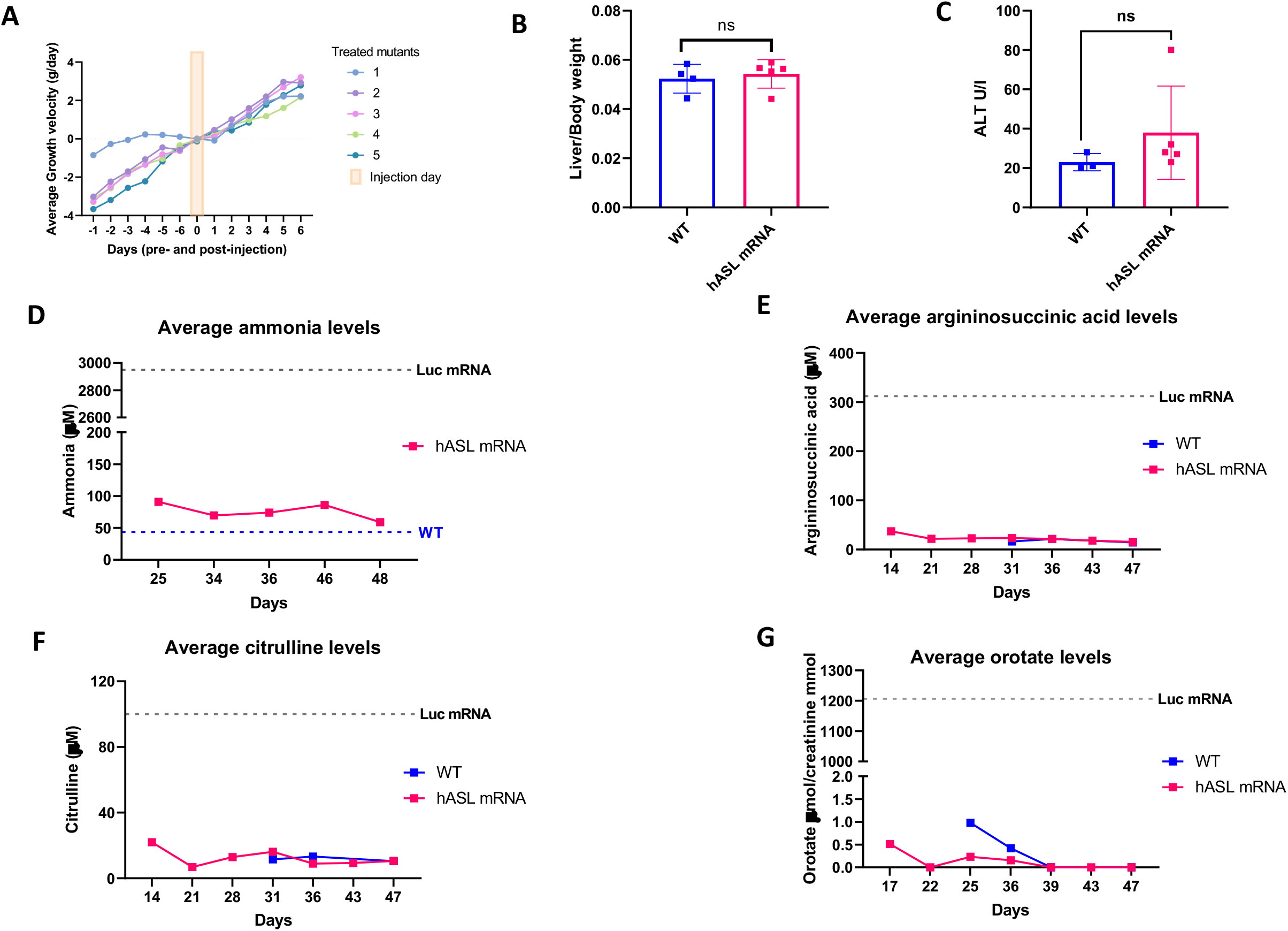
*hASL* mRNA therapy from birth corrects the phenotype of *Asl*^*Neo/Neo*^ mice. (A) Average growth velocity per *hASL* treated animals each day pre- and post-injection. (**B**) Liver to body weight ratio at harvest comparing WT against *hASL* mRNA treated *Asl*^*Neo/Neo*^ mice. (**C**) Plasma alanine aminotransferase (ALT) levels at harvest comparing WT against *hASL* mRNA treated *Asl*^*Neo/Neo*^ mice. Longitudinal (**D**) average plasma ammonia concentration, (**E**) argininosuccinic acid (**F)** citrulline from dried blood spots and (**G**) urine orotate from *hASL* mRNA treated *Asl*^*Neo/Neo*^ mice. (**B-C**) Unpaired two-tailed t-test per timepoint, ns= not significant. (**B-I**) WT n=3-4, *Asl*^*Neo/Neo*^ n=5 (**D**) Grey line indicates average levels from *Luc* mRNA treated *Asl*^*Neo/Neo*^ mice at harvest, blue line indicates average WT levels at harvest. (**E-G**) Grey line indicates average levels from *Luc* mRNA treated *Asl*^*Neo/Neo*^mice at harvest. (**A**) Graph shows mean per animal. (**B-C**) Graph show means +SD. (**D-G**) Graphs show mean per timepoint.

**Supplementary Figure 7.**
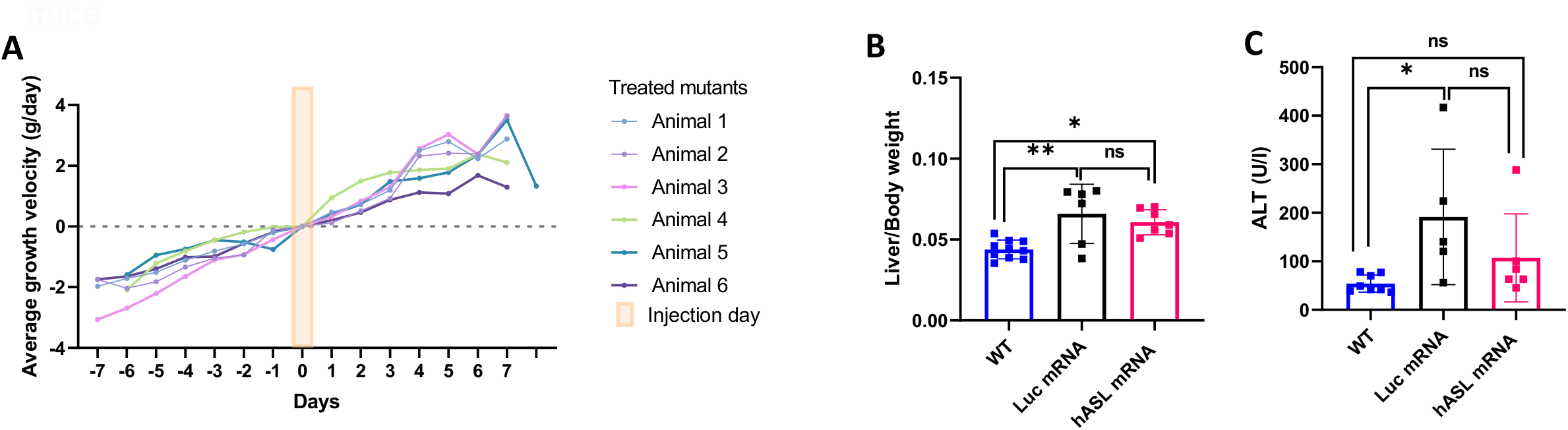
*hASL* mRNA therapy partially corrects the adult phenotype in *Asl*^*Neo/Neo*^ mice. (**A**) Average mean growth velocity per *hASL* treated animals each week pre- and post-injection. (**B**) Liver to body weight ratio at harvest comparing WT against hASL mRNA treated *Asl*^*Neo/Neo*^ mice showed no difference. (**C**) Plasma alanine aminotransferase (ALT) levels at harvest comparing WT against *hASL* mRNA treated *Asl*^*Neo/Neo*^ mice showed no difference. (**B-C**) One-way ANOVA with Tukey’s post-test, ns=not significant, *p<0.05, **p<0.01. (**B**) WT n=10, *Luc* mRNA n=6, *hASL* mRNA n=7 (**C**) WT n=8, *Luc* mRNA n=5, *hASL* mRNA n=8.

**Supplementary Figure 8.**
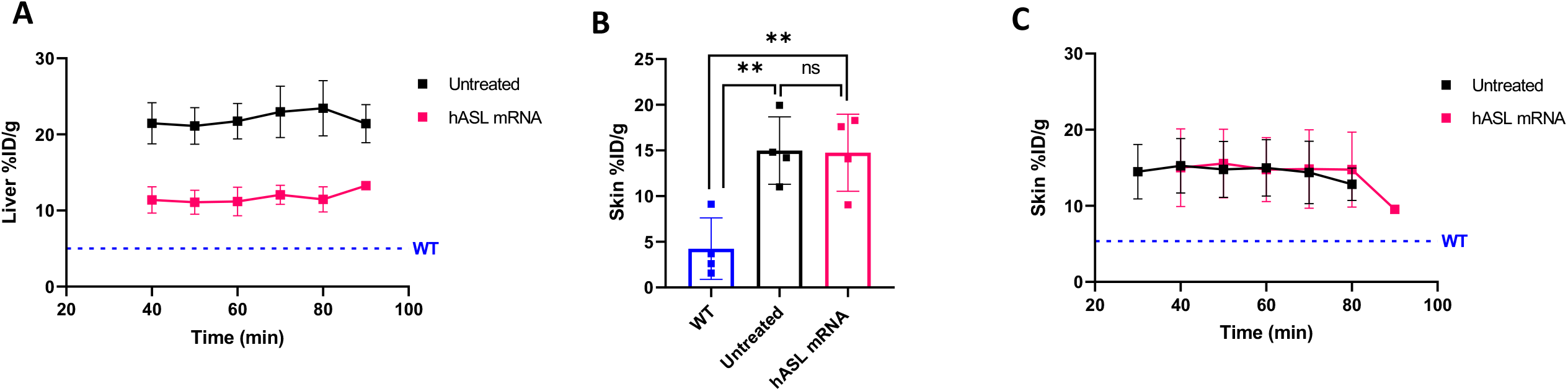
*hASL* mRNA therapy corrects the dysfunction of glutathione metabolism in *Asl*^*Neo/Neo*^ mice. Similarly, to Asl^Neo/Neo^ and WT mice, [^18^F] FSPG concentration (%ID/g) in mRNA treated *Asl*^*Neo/Neo*^ [^18^F] FSPG concentration (%ID/g) was more or less constant between 40 and 90 minutes post injection in liver (**A**) and skin (**C**). (**B**) [^18^F] FSPG retention in untreated *Asl*^*Neo/Neo*^ and *hASL* mRNA treated *Asl*^*Neo/Neo*^ mice was threefold higher than that of WT mice. Lack of efficacy of *hASL* mRNA in the skin is potentially due to *hASL* mRNA targeting the liver and not the skin.

## List of abbreviations

ASA: Argininosuccinic Aciduria
ASL: Argininosuccinate Lyase
ASLD: Argininosuccinate Lyase Deficiency
ASSD: Argininosuccinate Synthase Ddeficiency
CyS: Cysteine
CySS: Cystine
GSH: Glutathione
hASL: human Argininosuccinate Lyase
HcyS: Homocysteine
HcySS: Homocysteine
HILIC: Hydrophilic Interaction Liquid Chromatography
IP: Intraperitoneal
IV: Intravenous
IVIS: In vivo imaging system
LNP: Lipid nanoparticle
Luc: Luciferase
MDA: Malondialdehyde
NO: Nitric Oxide
OTCD: Ornithine Transcarbamylase Deficiency
PAS: Periodic Acid Schaff
PET: Positron Emission Tomography
ROI: Regions of Interest
TBARS: Thiobarbituric Acid reactive substances assay
WT: Wildtype
xCT: Cystine/Glutamate antiporter
[^18^F]FSPG: (4S)-4-(3-[^18^F]Fluoropropyl)-L-glutamate

## Notes

**Conflict of Interest:** The work was partly funded by Moderna Therapeutics.

**Funding:** This work was supported by funding from Moderna Therapeutics, the United Kingdom Medical Research Council Clinician Scientist Fellowship MR/T008024/1 (JB), London Advanced Therapy Confidence in Collaboration in Advanced Therapies award (2CiC017) (JB, THW), a Wellcome Trust Senior Fellowship 220221/Z/20/Z (THW), the CRUK City of London Centre Award C7893/A26233 (THW), and the NIHR Great Ormond Street Hospital Biomedical Research Centre (JB, PG, PBM, SE). The views expressed are those of the author(s) and not necessarily those of the NHS, the NIHR or the Department of Health.

**Author approvals:** All authors have seen and approved the manuscript. This manuscript has not been accepted or published elsewhere.

### Competing Interest Statement

The work was partly funded by Moderna Therapeutics.

## References

1. Summar ML, Koelker S, Freedenberg D, Le Mons C, Haberle J, Lee HS, et al. The incidence of urea cycle disorders. Mol Genet Metab. 2013;110(1-2):179–80.

2. Baruteau J, Diez-Fernandez C, Lerner S, Ranucci G, Gissen P, Dionisi-Vici C, et al. Argininosuccinic aciduria: Recent pathophysiological insights and therapeutic prospects. J Inherit Metab Dis. 2019;42(6):1147–61.

3. Erez A. Argininosuccinic aciduria: from a monogenic to a complex disorder. Genet Med. 2013;15(4):251–7.

4. Erez A, Nagamani SC, Shchelochkov OA, Premkumar MH, Campeau PM, Chen Y, et al. Requirement of argininosuccinate lyase for systemic nitric oxide production. Nat Med. 2011;17(12):1619–26.

5. Nagamani SC, Campeau PM, Shchelochkov OA, Premkumar MH, Guse K, Brunetti-Pierri N, et al. Nitric-oxide supplementation for treatment of long-term complications in argininosuccinic aciduria. Am J Hum Genet. 2012;90(5):836–46.

6. Nagamani SCS, Ali S, Izem R, Schady D, Masand P, Shneider BL, et al. Biomarkers for liver disease in urea cycle disorders. Mol Genet Metab. 2021;133(2):148–56.

7. Bigot A, Tchan MC, Thoreau B, Blasco H, Maillot F. Liver involvement in urea cycle disorders: a review of the literature. Journal of inherited metabolic disease. 2017;40(6):757–69.

8. Burrage LC, Madan S, Li X, Ali S, Mohammad M, Stroup BM, et al. Chronic liver disease and impaired hepatic glycogen metabolism in argininosuccinate lyase deficiency. JCI Insight. 2020;5(4).

9. Baruteau J, Jameson E, Morris AA, Chakrapani A, Santra S, Vijay S, et al. Expanding the phenotype in argininosuccinic aciduria: need for new therapies. J Inherit Metab Dis. 2017;40(3):357–68.

10. Tuchman M, Lee B, Lichter-Konecki U, Summar ML, Yudkoff M, Cederbaum SD, et al. Cross-sectional multicenter study of patients with urea cycle disorders in the United States. Mol Genet Metab. 2008;94(4):397–402.

11. Soria LR, Gurung S, De Sabbata G, Perocheau DP, De Angelis A, Bruno G, et al. Beclin-1-mediated activation of autophagy improves proximal and distal urea cycle disorders. EMBO Mol Med. 2021;13(2):e13158.

12. Marble M, McGoey RR, Mannick E, Keats B, Ng SS, Deputy S, et al. Living related liver transplant in a patient with argininosuccinic aciduria and cirrhosis: metabolic follow-up. J Pediatr Gastroenterol Nutr. 2008;46(4):453–6.

13. Zimmermann A, Bachmann C, Baumgartner R. Severe liver fibrosis in argininosuccinic aciduria. Arch Pathol Lab Med. 1986;110(2):136–40.

14. Seminotti B, da Silva JC, Ribeiro RT, Leipnitz G, Wajner M. Free Radical Scavengers Prevent Argininosuccinic Acid-Induced Oxidative Stress in the Brain of Developing Rats: a New Adjuvant Therapy for Argininosuccinate Lyase Deficiency? Mol Neurobiol. 2020;57(2):1233–44.

15. Jameson E BC JL, Vijay S, Morris AA. The Manchester experience with ASA patients. Journal of Inherited metabolic disorder; 2012.

16. Baruteau J, Perocheau DP, Hanley J, Lorvellec M, Rocha-Ferreira E, Karda R, et al. Argininosuccinic aciduria fosters neuronal nitrosative stress reversed by Asl gene transfer. Nat Commun. 2018;9(1):3505.

17. Touramanidou L PD, Gurung S, Cozmescu AC, Waddington SN, Counsell JR, Gissen P, Baruteau J. In vivo lentiviral gene therapy for argininosuccinic aciduria. European Society of Gene and Cell Therapy (ESGCT); Virtual Congress: Human Gene Therapy; 2021. p. A1–152.

18. Ashley SN, Nordin JML, Buza EL, Greig JA, Wilson JM. Adeno-associated viral gene therapy corrects a mouse model of argininosuccinic aciduria. Molecular genetics and metabolism. 2018;125(3):241–50.

19. Lu SC. Regulation of glutathione synthesis. Mol Aspects Med. 2009;30(1-2):42-59.

20. Mittra ES, Koglin N, Mosci C, Kumar M, Hoehne A, Keu KV, et al. Pilot Preclinical and Clinical Evaluation of (4S)-4-(3-[18F]Fluoropropyl)-L-Glutamate (18F-FSPG) for PET/CT Imaging of Intracranial Malignancies. PLoS One. 2016;11(2):e0148628.

21. Baek S, Choi CM, Ahn SH, Lee JW, Gong G, Ryu JS, et al. Exploratory clinical trial of (4S)-4-(3-[18F]fluoropropyl)-L-glutamate for imaging xC-transporter using positron emission tomography in patients with non-small cell lung or breast cancer. Clin Cancer Res. 2012;18(19):5427–37.

22. Kavanaugh G, Williams J, Morris AS, Nickels ML, Walker R, Koglin N, et al. Utility of [(18)F]FSPG PET to Image Hepatocellular Carcinoma: First Clinical Evaluation in a US Population. Mol Imaging Biol. 2016;18(6):924–34.

23. Park SY, Na SJ, Kumar M, Mosci C, Wardak M, Koglin N, et al. Clinical Evaluation of (4S)-4-(3-[(18)F]Fluoropropyl)-L-glutamate ((18)F-FSPG) for PET/CT Imaging in Patients with Newly Diagnosed and Recurrent Prostate Cancer. Clin Cancer Res. 2020;26(20):5380–7.

24. McCormick PN, Greenwood HE, Glaser M, Maddocks ODK, Gendron T, Sander K, et al. Assessment of Tumor Redox Status through (S)-4-(3-[(18)F]fluoropropyl)-L-Glutamic Acid PET Imaging of System xc (-) Activity. Cancer Res. 2019;79(4):853–63.

25. Greenwood HE, McCormick PN, Gendron T, Glaser M, Pereira R, Maddocks ODK, et al. Measurement of Tumor Antioxidant Capacity and Prediction of Chemotherapy Resistance in Preclinical Models of Ovarian Cancer by Positron Emission Tomography. Clin Cancer Res. 2019;25(8):2471–82.

26. Greenwood HE, Edwards R, Koglin N, Berndt M, Baark F, Kim J, et al. Radiotracer stereochemistry affects substrate affinity and kinetics for improved imaging of system xC (-) in tumors. Theranostics. 2022;12(4):1921–36.

27. Hoehne A, James ML, Alam IS, Ronald JA, Schneider B, D’Souza A, et al. [(18)F]FSPG-PET reveals increased cystine/glutamate antiporter (xc-) activity in a mouse model of multiple sclerosis. J Neuroinflammation. 2018;15(1):55.

28. Kho J, Tian X, Wong WT, Bertin T, Jiang MM, Chen S, et al. Argininosuccinate Lyase Deficiency Causes an Endothelial-Dependent Form of Hypertension. Am J Hum Genet. 2018;103(2):276–87.

29. Cortese-Krott MM, Koning A, Kuhnle GGC, Nagy P, Bianco CL, Pasch A, et al. The Reactive Species Interactome: Evolutionary Emergence, Biological Significance, and Opportunities for Redox Metabolomics and Personalized Medicine. Antioxid Redox Signal. 2017;27(10):684–712.

30. Abdulle AE, van Roon AM, Smit AJ, Pasch A, van Meurs M, Bootsma H, et al. Rapid free thiol rebound is a physiological response following cold-induced vasoconstriction in healthy humans, primary Raynaud and systemic sclerosis. Physiol Rep. 2019;7(6):e14017.

31. An D, Schneller JL, Frassetto A, Liang S, Zhu X, Park JS, et al. Systemic Messenger RNA Therapy as a Treatment for Methylmalonic Acidemia. Cell Rep. 2017;21(12):3548–58.

32. Jiang L, Park JS, Yin L, Laureano R, Jacquinet E, Yang J, et al. Dual mRNA therapy restores metabolic function in long-term studies in mice with propionic acidemia. Nat Commun. 2020;11(1):5339.

33. Ranucci G, Rigoldi M, Cotugno G, Bernabei SM, Liguori A, Gasperini S, et al. Chronic liver involvement in urea cycle disorders. J Inherit Metab Dis. 2019;42(6):1118–27.

34. Nagamani SC, Shchelochkov OA, Mullins MA, Carter S, Lanpher BC, Sun Q, et al. A randomized controlled trial to evaluate the effects of high-dose versus low-dose of arginine therapy on hepatic function tests in argininosuccinic aciduria. Mol Genet Metab. 2012;107(3):315–21.

35. Braissant O. Current concepts in the pathogenesis of urea cycle disorders. Mol Genet Metab. 2010;100 Suppl 1:S3–s12.

36. Parmeggiani B, Vargas CR. Oxidative stress in urea cycle disorders: Findings from clinical and basic research. Clin Chim Acta. 2018;477:121–6.

37. Aoyagi K, Nagase S, Gotoh M, Akiyama K, Satoh M, Hirayama A, et al. Role of reactive oxygen and argininosuccinate in guanidinosuccinate synthesis in isolated rat hepatocytes. Enzyme Protein. 1996;49(4):205–11.

38. Wink DA, Miranda KM, Espey MG, Pluta RM, Hewett SJ, Colton C, et al. Mechanisms of the antioxidant effects of nitric oxide. Antioxid Redox Signal. 2001;3(2):203–13.

39. Feelisch M, Cortese-Krott MM, Santolini J, Wootton SA, Jackson AA. Systems redox biology in health and disease. EXCLI J. 2022;21:623–46.

40. Meister A. Glutathione metabolism. Methods Enzymol. 1995;251:3–7.

41. Seelig GF, Simondsen RP, Meister A. Reversible dissociation of gamma-glutamylcysteine synthetase into two subunits. J Biol Chem. 1984;259(15):9345–7.

42. Chen Y, Shertzer HG, Schneider SN, Nebert DW, Dalton TP. Glutamate cysteine ligase catalysis: dependence on ATP and modifier subunit for regulation of tissue glutathione levels. J Biol Chem. 2005;280(40):33766–74.

43. Jin Z, Kho J, Dawson B, Jiang MM, Chen Y, Ali S, et al. Nitric oxide modulates bone anabolism through regulation of osteoblast glycolysis and differentiation. The Journal of clinical investigation. 2021;131(5).

44. Kensler TW, Wakabayashi N, Biswal S. Cell survival responses to environmental stresses via the Keap1-Nrf2-ARE pathway. Annu Rev Pharmacol Toxicol. 2007;47:89–116.

45. Vairetti M, Di Pasqua LG, Cagna M, Richelmi P, Ferrigno A, Berardo C. Changes in Glutathione Content in Liver Diseases: An Update. Antioxidants (Basel). 2021;10(3).

46. García-Tevijano ER, Berasain C, Rodríguez JA, Corrales FJ, Arias R, Martín-Duce A, et al. Hyperhomocysteinemia in liver cirrhosis: mechanisms and role in vascular and hepatic fibrosis. Hypertension. 2001;38(5):1217–21.

47. Snyderman SE. Liver failure and neurologic disease in a patient with homocystinuria. Mol Genet Metab. 2006;87(3):210–2.

48. Maclean KN, Sikora J, Kožich V, Jiang H, Greiner LS, Kraus E, et al. Cystathionine beta-synthase null homocystinuric mice fail to exhibit altered hemostasis or lowering of plasma homocysteine in response to betaine treatment. Mol Genet Metab. 2010;101(2-3):163–71.

49. Majtan T, Hůlková H, Park I, Krijt J, Kožich V, Bublil EM, et al. Enzyme replacement prevents neonatal death, liver damage, and osteoporosis in murine homocystinuria. Faseb j. 2017;31(12):5495–506.

50. Liu X, Zhang Y, Zhuang L, Olszewski K, Gan B. NADPH debt drives redox bankruptcy: SLC7A11/xCT-mediated cystine uptake as a double-edged sword in cellular redox regulation. Genes Dis. 2021;8(6):731–45.

51. Liu J, Xia X, Huang P. xCT: A Critical Molecule That Links Cancer Metabolism to Redox Signaling. Mol Ther. 2020;28(11):2358–66.

52. Hambraeus L, Hardell LI, Westphal O, Lorentsson R, Hjorth G. Argininosuccinic aciduria. Report of three cases and the effect of high and reduced protein intake on the clinical state. Acta Paediatr Scand. 1974;63(4):525–36.

53. Schutgens RB, Beemer FA, Tegelaers WH, de Groot WP. Mild variant of argininosuccinic aciduria. J Inherit Metab Dis. 1980;2(1):13–4.

54. Baruteau J, Waddington SN, Alexander IE, Gissen P. Gene therapy for monogenic liver diseases: clinical successes, current challenges and future prospects. Journal of inherited metabolic disease. 2017;40(4):497–517.

55. Wilson JM, Flotte TR. Moving Forward After Two Deaths in a Gene Therapy Trial of Myotubular Myopathy. Human gene therapy. 2020;31(13-14):695–6.

56. Philippidis A. Fourth Boy Dies in Clinical Trial of Astellas’ AT132. Human gene therapy. 2021;32(19-20):1008–10.

57. Guillou J, de Pellegars A, Porcheret F, Fremeaux-Bacchi V, Allain-Launay E, Debord C, et al. Fatal thrombotic microangiopathy case following adeno-associated viral SMN gene therapy. Blood Adv. 2022;6(14):4266–70.

58. Sahin U, Karikó K, Türeci Ö. mRNA-based therapeutics--developing a new class of drugs. Nat Rev Drug Discov. 2014;13(10):759–80.

59. Berraondo P, Martini PGV, Avila MA, Fontanellas A. Messenger RNA therapy for rare genetic metabolic diseases. Gut. 2019;68(7):1323–30.

60. Córdoba KM, Jericó D, Sampedro A, Jiang L, Iraburu MJ, Martini PGV, et al. Messenger RNA as a personalized therapy: The moment of truth for rare metabolic diseases. Int Rev Cell Mol Biol. 2022;372:55–96.

61. Hou X, Zaks T, Langer R, Dong Y. Lipid nanoparticles for mRNA delivery. Nat Rev Mater. 2021;6(12):1078–94.

62. Damase TR, Sukhovershin R, Boada C, Taraballi F, Pettigrew RI, Cooke JP. The Limitless Future of RNA Therapeutics. Front Bioeng Biotechnol. 2021;9:628137.

63. An D, Frassetto A, Jacquinet E, Eybye M, Milano J, DeAntonis C, et al. Long-term efficacy and safety of mRNA therapy in two murine models of methylmalonic acidemia. EBioMedicine. 2019;45:519–28.

64. Yu H, Brewer E, Shields M, Crowder M, Sacchetti C, Soontornniyomkij B, et al. Restoring ornithine transcarbamylase (OTC) activity in an OTC-deficient mouse model using LUNAR-OTC mRNA. Clinical and Translational Discovery. 2022;2(2):e33.

65. Khoja S, Liu XB, Truong B, Nitzahn M, Lambert J, Eliav A, et al. Intermittent lipid nanoparticle mRNA administration prevents cortical dysmyelination associated with arginase deficiency. Mol Ther Nucleic Acids. 2022;28:859–74.

66. Truong B, Allegri G, Liu XB, Burke KE, Zhu X, Cederbaum SD, et al. Lipid nanoparticle-targeted mRNA therapy as a treatment for the inherited metabolic liver disorder arginase deficiency. Proc Natl Acad Sci U S A. 2019;116(42):21150–9.

67. Nelson J, Sorensen EW, Mintri S, Rabideau AE, Zheng W, Besin G, et al. Impact of mRNA chemistry and manufacturing process on innate immune activation. Sci Adv. 2020;6(26):eaaz6893.

68. Edwards R, Greenwood HE, McRobbie G, Khan I, Witney TH. Robust and Facile Automated Radiosynthesis of [(18)F]FSPG on the GE FASTlab. Mol Imaging Biol. 2021;23(6):854–64.

69. Greenwood HE, Nyitrai Z, Mocsai G, Hobor S, Witney TH. High-Throughput PET/CT Imaging Using a Multiple-Mouse Imaging System. J Nucl Med. 2020;61(2):292–7.

70. Prinsen H, Schiebergen-Bronkhorst BGM, Roeleveld MW, Jans JJM, de Sain-van der Velden MGM, Visser G, et al. Rapid quantification of underivatized amino acids in plasma by hydrophilic interaction liquid chromatography (HILIC) coupled with tandem mass-spectrometry. J Inherit Metab Dis. 2016;39(5):651–60.

71. Janero DR, Bryan NS, Saijo F, Dhawan V, Schwalb DJ, Warren MC, et al. Differential nitros(yl)ation of blood and tissue constituents during glyceryl trinitrate biotransformation in vivo. Proceedings of the National Academy of Sciences of the United States of America. 2004;101(48):16958–63.

72. McKenna HT, O’Brien KA, Fernandez BO, Minnion M, Tod A, McNally BD, et al. Divergent trajectories of cellular bioenergetics, intermediary metabolism and systemic redox status in survivors and non-survivors of critical illness. Redox Biol. 2021;41:101907.

73. Sutton TR, Minnion M, Barbarino F, Koster G, Fernandez BO, Cumpstey AF, et al. A robust and versatile mass spectrometry platform for comprehensive assessment of the thiol redox metabolome. Redox Biol. 2018;16:359–80.

74. Kok CY, Cunningham SC, Carpenter KH, Dane AP, Siew SM, Logan GJ, et al. Adeno-associated virus-mediated rescue of neonatal lethality in argininosuccinate synthetase-deficient mice. Molecular therapy : the journal of the American Society of Gene Therapy. 2013;21(10):1823–31.

75. Love MI, Huber W, Anders S. Moderated estimation of fold change and dispersion for RNA-seq data with DESeq2. Genome Biol. 2014;15(12):550.

76. Wickham H. ggplot2: Elegant Graphics for Data Analysis. Springer-Verlag New York ISBN 978-3-319-24277-4. 2016;https://ggplot2.tidyverse.org.

